# Axiomatic Community Ecology, Topology, and Dynamic Distance

**DOI:** 10.1101/2025.08.21.671550

**Authors:** Ned J.H. Wontner, Matthew Spencer

## Abstract

The super-organismal view of ecosystems has largely been superseded by the individualistic view. One consequence of the dominance of the individualistic view is that many modern ecologists treat ecosystems as nothing more than vectors of relative abundances, ignoring the potentially important idea that an understanding of ecosystems should be based on dynamics rather than abundances or species identities. We develop a mathematical framework in which we compare dynamical properties of ecosystems with different sets of species, using ideas from functional analysis, metric spaces, and topology. We give two proof-of-principle applications of our framework to marine sessile communities and to a large database of ecosystem models. We show that under a set of biologically-motivated axioms designed to capture the properties of predator-prey systems, there is only one natural kind of ecosystem.

## 1 Introduction

The study of ecosystems by English-speaking ecologists has included at least two main lines of thought. One tradition (the super-organismal view) holds that ecosystems are well-defined units, with properties whose study requires a different approach from that needed at lower levels of organisation (Clements and Shelford, 1939, pp. 20-25). An opposing tradition (the individualistic view) holds that ecosystems are not well-defined units (Whittaker, 1975, pp. 113-116) and that their properties can be understood at the level of the species they contain (Gleason, 1926). Note that throughout we will use the term “ecosystem” to denote a set of species and the nature of their interactions with each other (in a sense that we make precise below). Historically, other terms including “association”, “formation”, “biome” and “community” would all have overlapped with our use of “ecosystem”, but with different implications about the level of distinct organisation. However, with the perspective of almost 100 years, the differences between these terms do not seem as important as they may have done in the mid 20th century, and the term “ecosystem” originated in an attempt to synthesize aspects of both traditions (Tansley, 1935). Our perspective is also mainly informed by North American and British ecology, although there were clear links to European plant ecology (Nicolson, 1996) and biogeochemistry (Pireddu, 2023). It is now generally believed that the super-organismal view of ecosystems is incorrect (Hagen, 1992, chapter 3), and descendant systems-level approaches (Odum, 1983, chapters 2, 21, 22) are out of fashion. One consequence of the dominance of the individualistic view is that potentially important ideas about the nature of ecosystems that originated with the super-organismal view are rarely considered when comparing and classifying ecosystems. In particular, Clements argued that the abundance of each species is not a key feature of ecosystems (Clements and Shelford, 1939, p. 249), that the understanding of dynamical processes is fundamental to the understanding of ecosystems (Clements and Shelford, 1939, p. 248), and that differences in life forms are more fundamental than differences in species identities (Clements and Shelford, 1939, p. 230). It seems likely that focusing on dynamics rather than abundances or species identities remains relevant to modern ecology.

Modern ecologists working in the mainstream of the subject, and especially in applied areas, often ignore the possibility of organisation at the ecosystem level and treat an ecosystem as nothing more than a vector of relative abundances. A typical approach is to compute some measure of dissimilarity in relative abundances, such as Bray-Curtis distance, and use non-metric multidimensional scaling and permutational analysis of variance to look for patterns of difference (e.g. Anderson, 2001). The stated aim of this approach is “to describe and interpret the structure of data sets by combining a variety of numerical approaches” (Legendre and Legendre, 2012, p. xiv). However, this approach makes little use of ecological theory, which limits its ability to provide explanations. In contrast, there are ways to compare ecosystems that make use of ecosystem-level properties. For example, Ulanowicz (1986) is a serious attempt to describe the structure of ecological flow networks using information theory, with much clearer mathematical and ecological foundations than Odum (1983). Ulanowicz et al. (2014) used data on amounts of energy flowing between each species, and ideas from community ecology such as the instability of highly connected networks, to suggest limits on the complexity of ecosystems. A systematic review of related work on ecological network analysis (Borrett et al., 2018) suggests the potential for broad applicability in fields such as food web status and effects of toxins. Although the focus on energy flow has attracted controversy from population biologists for many decades (e.g. Slobodkin, 1960), there is scope for further work on ecosystem comparison based on processes rather than just abundances, and on well-established theory about community dynamics.

The axiomatic approach to community ecology began with the mathematician Kolmogorov, who made an early and surprisingly non-technical contribution to the development of predator-prey models (Kolmogorov, 1936). Rather than specifying a particular model for a predator-prey system (such as a Lotka-Volterra model), Kolmogorov chose a set of axioms designed to capture the essential properties of all predator-prey models, for example that the proportional population growth rate of the prey species decreases as the number of predators increases. With these axioms, he proved non-trivial results such as the range of possible long-term dynamical behaviours for a predator-prey system. This approach has since been fruitful in mathematical biology (Sigmund, 2007). For example, with a very small number of axioms about the forms of differential equations describing competitive systems, it is possible to prove general results about how the complexity of possible dynamics increases with the number of competing species (Smale, 1976). Axiomatic approaches have had less impact in ecology than in mathematical biology, perhaps because a higher level of mathematical sophistication is needed to study axioms about dynamics than to study particular models. However, there have been exceptions. For example, Hutchinson (1978, pp. 1-4) used axioms such as the postulate of parenthood (that every living organism arises from at least one parent of the same kind) and the postulate of an upper limit on the number of organisms that can live in a finite space to justify the logistic model at a level accessible to undergraduates. Hutchinson’s mathematical approach was based on Lotka (1956, pp. 64-65), but he gave much more detail on the biological reasoning underlying the axioms, even though his focus was on one particular model satisfying those axioms. More recently, an axiomatic approach has been used to identify the kinds of functions describing interspecific interactions that do not lead to inconsistencies if populations are arbitrarily split into clones (Ansmann and Bollenbach, 2021). It is likely that axiomatic community ecology can provide stronger links between the principles of population biology and methods for comparing and classifying ecosystems.

Here, we develop a mathematical framework which compares ecosystem models through their dynamical properties at the level of pairwise interactions, rather than abundances or species identities. We apply this framework to two data sets: an experimental study of marine sessile communities and a large database of ecosystem models. We show that under a reasonable axiomatic refinement of the framework, there is a broad class of measurements of dynamical properties for which there is just one kind of ecosystem.

## 2 Outline of Approach

Here, we give an overview of the rest of the paper. Throughout, we have given definitions and theorems in the main text, but proofs and other technical details are in appendices.

In Section 3, we recall some standard definitions from topology and metric spaces. We then introduce some definitions from functional analysis, in particular distances between real functions, distances between matrices, and thus distances between matrices of real functions. In our case, these matrices represent pairwise interactions between species, and distances between them are used to measure dynamical distance between ecosystems. To deal with the situation where different ecosystems have different sets of species, we introduce a non-standard definition of a pseudodistance, where we minimise the distance over all the possible ways to match each species in the smaller ecosystem with exactly one species in the larger ecosystem. Figure 1 shows the main steps in our approach. The approach is general, although some choices must be made about details in particular approaches. We give two examples below (Sections 4 and 5), where we show as proof of concept that we can identify biologically meaningful patterns in the dynamics of two sets of ecosystem models.

**Figure 1:**
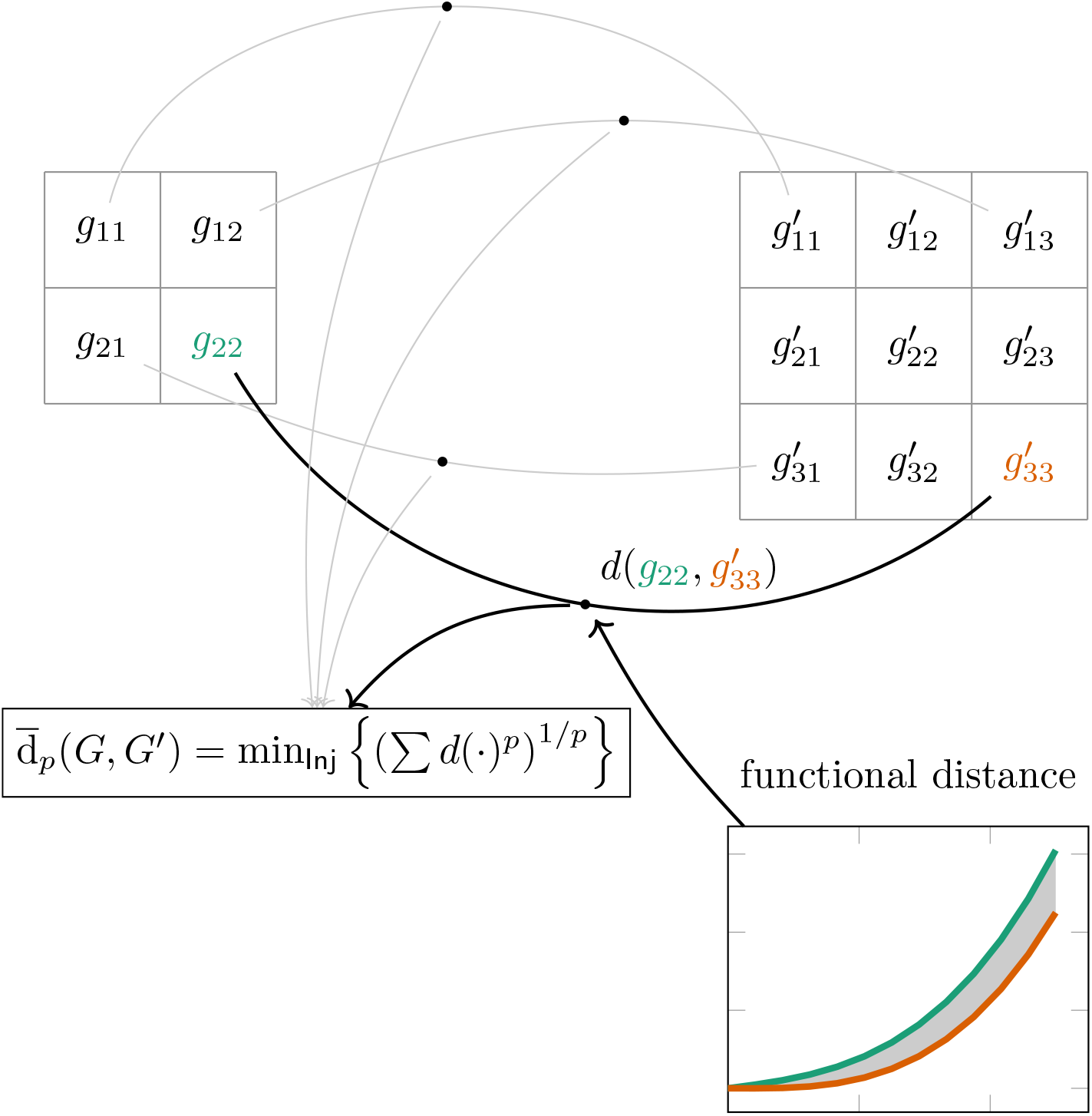
Outline of our approach to calculating pseudodistances between ecosystems. Given two matrices *G, G*′ of functions representing pairwise interactions between species, we match each species in the smaller ecosystem with exactly one species in the larger ecosystem. For example, here species 1 in the 2-species ecosystem is matched with species 1 in the 3-species ecosystem, and species 2 in the 2-species ecosystem is matched with species 3 in the 3-species ecosystem. We then compute functional distances *d* between matching functions (e.g. *g*_22_ in *G*, green, and 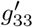 in *G*′, orange) using the 1*/p* power of the integral of some power *p* of the absolute difference between the functions (lower right, grey shaded area). For example, *d*(*g*_22_, 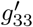) on the figure is one of the functional distances *d*(*p, i, j, σ*) of Definition 3.3. We sum the *p*th powers of these functional distances over all matching pairs and take the 1*/p* power of the sum. We minimize this sum over all the possible ways to match each species in the smaller ecosystem with exactly one species in the larger ecosystem (the set of injective mappings, denoted Inj). This gives the pseudodistance 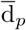(*G, G*′).

In Section 6, we introduce a collection of biologically-motivated axioms which characterise a formal mathematical representation of ecosystems that do not have immigration or emigration. We can use the metric-like functions introduced above to show that ecosystems satisfying these axioms form a space which resembles a topology (informally, one in which points can be near to or far from each other, even though distance may not be numerically measurable). When we restrict to ecosystems with only production and predation, a minor modification of a well-known theorem shows that this space is connected (informally, cannot be split naturally into two or more parts). A standard theorem then shows that any continuous function (informally, one that can be drawn without lifting the pen from the paper) on the space has a connected image. In other words, any natural measurement gives just one kind of ecosystem (Figure 2).

**Figure 2:**
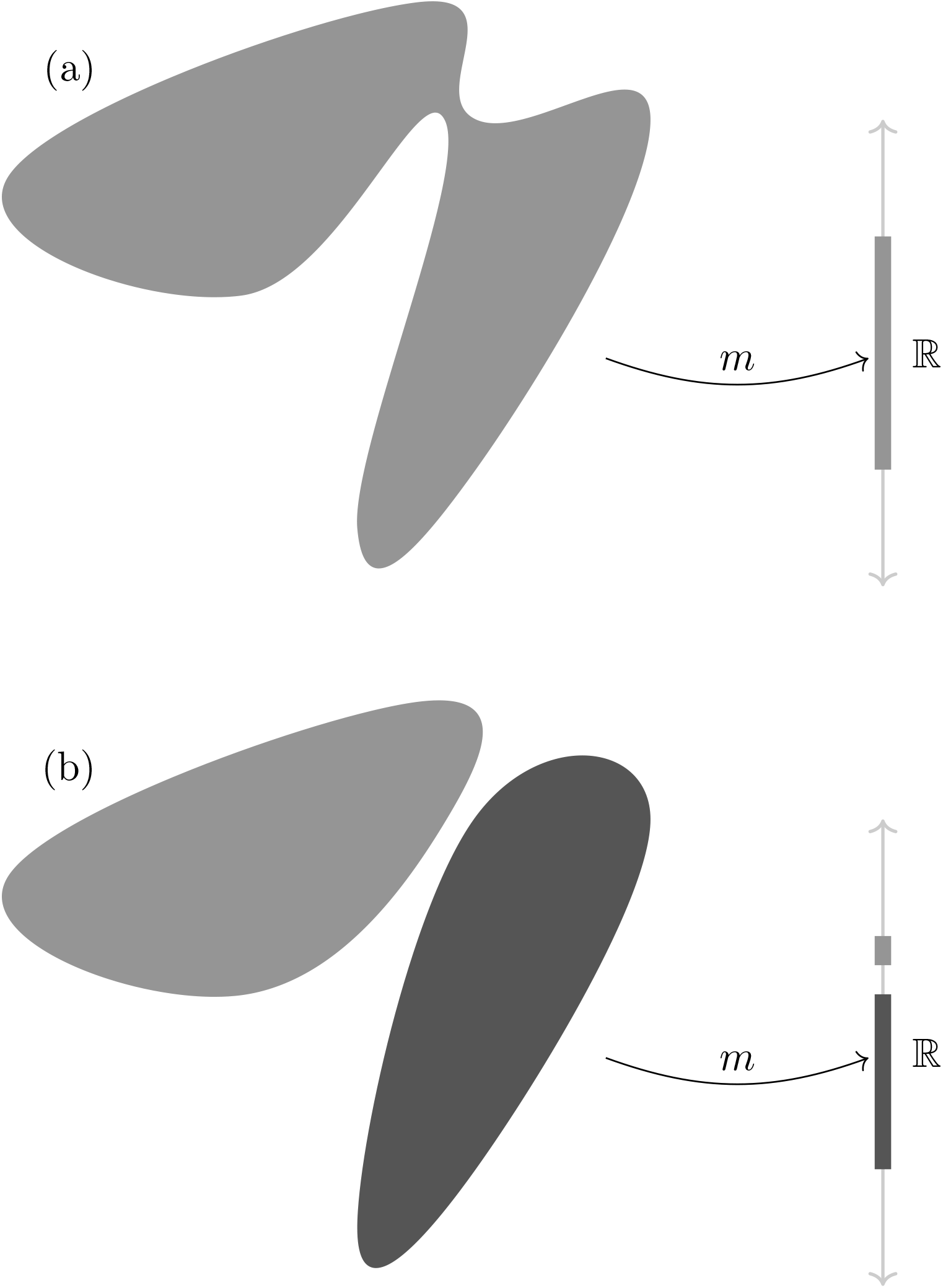
Connected spaces and natural measurements. In (a), the grey shape represents a connected space, such as the space of ecosystems satisfying our axioms. A natural measurement *m* is a continuous real-valued function on this space. The image of the space under this function is connected (an interval of the real line ℝ), so that there is just one kind of ecosystem with respect to this natural mapping. In contrast, if the space is not connected (b), its image under a natural measurement may not be connected, and there may therefore be more than one kind of ecosystem with respect to this natural mapping.

## 3 Pseudodistances Between Ecosystems

Here, we specify the mathematical notions which we use to measure the dynamical distance between models of ecosystems. We do this in two steps. Firstly, we recall two classical notions, namely metrics and topologies, which mathematise the informal notions of distance and space, respectively. Secondly, we overview the relevant generalisation of metrics, called pseudometrics, which allow distinct points in a space to be at distance 0 from one another. We hence define pseudometrics and topology-like families on models of ecosystems (see also Section 6). Proofs are found in Appendix E.

Throughout, we use the following notation. We write *X, Y*, … for sets, *X × Y* := *{*(*x, y*) : *x* ∈ *X, y* ∈ *Y}, P*(*X*) := *{Y* ⊆ *X}*, and ∅ for the empty set. We let ℕ be the set of natural numbers 0, 1, 2, …, whose members we denote by *k, l, m, n*, We let *X × X* =: *X*^2^ and likewise *X*^*n*^ consists of the vectors **x** with *n* coordinates, which we write *x*_*n*_, hence we write **x** = (*x*_*m*_)_*m*∈*n*_. Similarly, we write *X*^*k×k*^ for the set of (square) matrices with width *k*, where each value in the matrix is an element of *X*, we write *G, G*′, ∈ *X*^*k×k*^ for the matrices, and denote their (*i, j*)^*th*^ entries by *g*_*i,j*_, hence we write *G* = (*g*_*i,j*_)_*i,j*≤*k*_.

We write ℝ for the set of real numbers, whose members we denote by *r, s*, If *L* is an order and *x* ∈ *L*, we write *L*_≥*x*_ := *{y* ∈ *L* : *y* ≥ *x}* and likewise *L*_*>x*_ := *{y* ∈ *L* : *y > x}* (e.g. ℝ_≥0_ and ℕ_*>*0_); and write *I* ⊆ ℝ for an interval in ℝ. We write *X*^*Y*^ for the set of functions from *Y* to *X*, and denote the functions by *e, f*,; if *f* : *X ×X* → ℝ_≥0_, we write *B*_*f*_ (*x, r*) := *{y* ∈ *X* : *f* (*x, y*) *< r}* which we call the *open ball of radius r around a point x*. Let *C*^∞^(*X*) ⊆ (ℝ_≥0_)^*X*^ be the set of positive real-valued function of *X* which are infinitely differentiable. Let 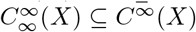 such that, if 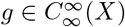, then for all *p* ∈ ℝ_≥1_, |*g*|^*p*^ is Lebesgue-measurable (see Tao, 2009, for more on measurability).

In general, we write *x*_*i*_(*t*) for species abundance, and *g*_*i,j*_(*x*_*i*_, *x*_*j*_) for the pairwise effects of species *i* on species *j*.

### 3.1 Metrics and Topologies

Our standard definitions follow Sutherland (2009). Let *X* be a set. A function *d* : *X × X* → ℝ_≥0_ is called a *metric* if, for all *x, y, z* ∈ *X*, the following hold:

M1. *d*(*x, y*) = 0 if and only if *x* = *y*,

M2. *d*(*x, y*) = *d*(*y, x*), and

M3. *d*(*x, y*) + *d*(*y, z*) ≥ *d*(*x, z*).

A metric formalises the notion of distance. Slightly more generally, a *topology* formalises the informal notion of a space (i.e. where points can be near or far, but the distance might not be numerically measurable). We say that *τ* ⊆ *p*(*X*) is a topology on *X* if:

T1. ∅ ∈ *τ*,

T2. if *Y, Z* ∈ *τ* then *Y* ∩ *Z* ∈ *τ* (closure under binary intersections), and

T3. if *Y*_*i*_ ∈ *τ* for all *i* ∈ *I*, then ⋃ _*i*∈*I*_ *Y*_*i*_ ∈ *τ* (closure under arbitrary unions).

We call *Y* ∈ *τ* an *open set* of *τ*. Informally, *x* and *y* are close together if a typical open set containing *x* also contains *y*. Open balls form a topology: let *d* be a metric on *X*. Then {U_*i*∈*I*_ *B*_*d*_(*x*_*i*_, *r*_*i*_) : *i* ∈ *I, x*_*i*_ ∈ *X, r*_*i*_ ∈ ℝ_≥0_*}* is a topology on *X*.

We use certain generalisations of connectedness and continuity. Informally, a space is connected if it cannot be split into two natural (i.e. open) parts, and a function is continuous if it can be drawn without taking the pen off the paper (so, for example, a sine wave is continuous, but a step function is not).

We use slight generalisations of these, as our collection of subsets need not be closed under intersections, so need not be topologies (cf. (Wontner, 2023, Chapter 2)). Let *τ* and *σ* be families of subsets of *X* and *Y* respectively. We say that *A* ⊆*X* is *(τ-)connected* if there is no pair of disjoint sets *U, V* ∈*τ* such that *A* ⊆*U*∪ *V*. If *f* : *X*→ *Y* is such that for any *A*∈ *σ, f*^−1^[*A*] ∈ *τ*, then we say that *f* is ((*τ, σ*)*-)continuous*. Examples includes *x*^2^ : ℝ →ℝ, ln(*x*) : ℝ_*>*0_ →ℝ, the composition of two continuous functions, and so on. The following is a mild generalisation of a standard fact:

#### Proposition 3.1.

(Wontner, 2023, Theorem 6.1.1) Let *τ* and *σ* be families of subsets of *X* and *Y* respectively. Suppose *f* : *X* → *Y* is continuous. If *A* ⊆ *X* is (*τ*-)connected then *f* (*A*) is (*σ*-)connected.

We now sketch particular metrics. A standard check shows that all of the following are indeed metrics. A simple example, on ℝ, is the size of the difference, i.e. *d*(*r, s*) := |*r* − *s*|. This is the everyday notion of distanceJon ℝ. If *n* ∈ ℕ_*>*1_, then we typically use the Euclidean distance on ℝ^*n*^: if **x, y** ∈ ℝ^*n*^, then 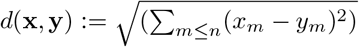 (also called the *l*_2_*-metric*). More generally, we have the *l*_*p*_*-metric*: let *p* ∈ ℝ be such that *p* ≥ 1. Fix an *n* ∈ ℕ_≥1_. If **x, y** ∈ ℝ^*n*^, we define 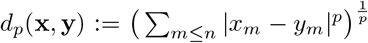 Similarly, we let *d*_∞_(**x, y**) := max*{*|*x*_*m*_ − *y*_*m*_| : *m* ≤ *n}*.

The previous metrics are on ℝ^*n*^ for *n* ∈ ℕ. We define metrics on matrices of real numbers like so: let *G, G*′ ⊆ ℝ^*k×k*^ with *G* = (*g* _*i,j*_)_*i,j*≤*k*_ and 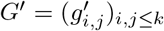. Then 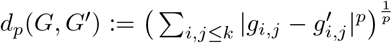, and likewise for *d*_∞_(*G, G*′).

We are interested in metrics on sets of matrices of functions. Such matrices are defined in Magnus and Neudecker (2019, chapter 5, section 15) and have applications in fields including statistics and econometrics. In our case it is most natural to think of them as functions *G* : (ℝ^2^→ℝ)^*n×n*^ representing pairwise effects of abundances on proportional population growth rates for sets of *n* species. We define metrics on real-valued functions corresponding to those of *l*_*p*_, these are the *L*_*p*_*-metrics* ((Sutherland, 2009, pages 45-47)). Let *S* ⊆ ℝ^*n*^ for some *n* ∈ ℕ. Let 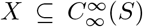 (i.e. *X* is a set of integrable real-valued functions on *S*). Let *p* ∈ ℝ_≥1_. Let *f, g* ∈ *X*. Then 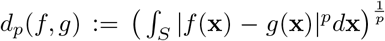. Similarly, we define *d*_∞_(*f, g*) := max_**x**∈*S*_ |*f* (**x**) −*g*(**x**)|. The corresponding *d*_*p*_ and *d*_∞_ metrics are defined for functions *f, g* : ℝ^*n*^ → ℝ, for example 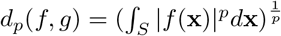.

We can define a metric on sets of vectors or matrices of real-valued functions, just as we did on sets of real-valued vectors or matrices. For simplicity, we write down the matrix case:

#### Definition 3.2.

Let *G* = (*g*_*i,j*_)_*i,j*≤*k*_ and 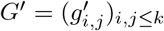 where each 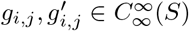 for some *S* ⊆ ℝ^*n*^. We define a metric on (ℝ^*S*^)^*k×k*^ by first using the (ℝ^*n*^ version of the) *L*_*p*_-metric, then using the *l*_*q*_-metric: 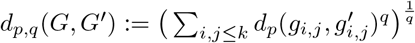. Similar versions using the *d*_∞_ metrics can also be stated.

### 3.2 Pseudometrics

When measuring the distances between the dynamics of the ecosystems, the labelling of the species is arbitrary: it should not matter if a species is labelled 1, or labelled 3141. However, in Definition 3.2, relabelling the species (i.e. permuting the labels) may yield a matrix with non-zero distance from the original matrix. So, we use a generalisation of metrics, called pseudometrics (cf. (Engelking, 1989, page 248)), which is permutation-independent.

Let *X* be a set. A function 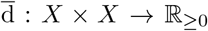 is called a *pseudometric* if, for all *x, y, z ∈ X*, the following hold:

P1. 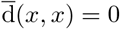,

P2. 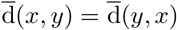, and

P3. 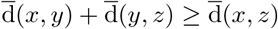.

Only the first conditions differs between metrics and pseudometrics: a pseudometric allows distinct points to have ‘distance’ 0 from one another. Like a metric, we can use a pseudometric 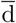 on a set *X* to form a topology (letting *A* be open if and only if for all *x* ∈*A*, there is an *r >* 0 such that 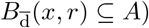.

If we minimise over the possible permutations of the labels in a square matrix, the definitions of matrix metrics from Section 3.1 yields pseudometrics. For any set *X*, let Perm(*X*) be the set of permutations of *X*, i.e. the set of bijections *σ* : *X* → *X*. If *k* ∈ ℕ, we write *σ* ∈ Perm(*k*) for a bijection *σ* : *{*1, …, *k}* → {1, …, *k*}. If *G*∈*Y* ^*k×k*^, we write *σ*(*G*) for the matrix (*g*_*σ*(*i*),*σ*(*j*)_)_*i,j*≤*k*_.

The simple case of a pseudometric generated by permuting a (square) matrix metric is when the entries in the matrix are real numbers: let *G, G*′ ⊆ ℝ^*k×k*^. Then:

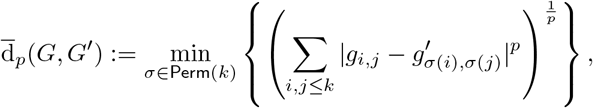

and likewise for 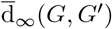.

The case with matrices of real-valued functions is similar.

#### Definition 3.3.

Let *k* ∈ ℕ. Let *G* = (*g*_*i,j*_)_*i,j*≤*k*_ where each 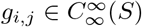 for some *S* ⊆ ℝ^*n*^ with *n* ∈ ℕ.

Let *p, q* ∈ ℝ_≥1_. Then, as a shorthand, we write

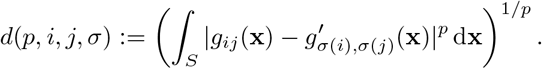

We define a function on 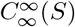 by first using the *L*_*p*_-metric, then using the *l*_*q*_-metric:

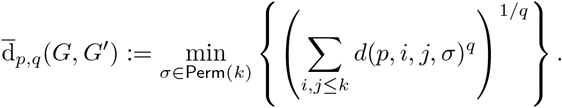

Similar versions using the ∞-metrics can also be stated. We use the following shorthand:

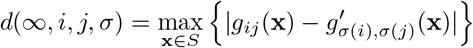

We then define the following function, which we will shortly show is a pseudometric:

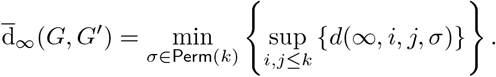

When *G, G*′ are clear, we write 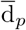 for 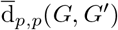 and 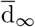 for 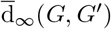 (e.g. in Sections 4 and 5).

This is how we pseudometrise the space of ecosystems: we identify (a model of) an ecosystem by a certain matrix of real-valued functions representing a certain way of capturing the dynamical behaviour of the ecosystems. Ours will be 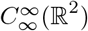 functions; in Section 6, these represent the proportional effect of one species on the rate of change of abundance on another species, etc. Variants of this pseudometric can be found in Appendix D.

If *k* is fixed in Definition 3.3, e.g. two ecosystems have the same number of species, then 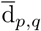 is a pseudometric:

#### Theorem 3.4.

Let *k* ∈ ℕ, and *p* = ∞ or *p* ∈ ℝ_≥1_. Then 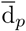 on ℝ^*k×k*^ is a pseudometric. Moreover, if *S* ⊆ ℝ^*n*^ is bounded, and *q* = ∞ or *q* ∈ ℝ_≥1_, then d_*p,q*_ is a pseudometric on 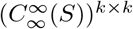.

Definition 3.3 allows us to measure the distance between matrices (e.g. ecosystems) when both have the same number of labels (so same number of species). We also want to be able to compare ecosystems with different numbers of labels. We cannot straightforwardly use the Definition 3.3. Instead, we vary Definition 3.3 in two ways. First, we use injections rather than permutation (as the two ecosystems may have different number of species). Secondly, we need to be slightly careful about dimensionality: in some cases (e.g. when certain variables are not dimensionless), we need to ensure that the defined function has the same dimensions for any pair of ecosystems, which we achieve through a normalising function (see Section 5.2 for an example).

#### Definition 3.5.

Let *S* ⊆ ℝ for some *n* ∈ ℕ and *l, k* ∈ ℕ where *l* ≤ *k* ≤ *n*. Let 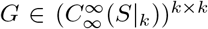 and 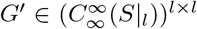, where *S*|_*k*_ is the restriction of *S* to ℝ^*k*^, and so too for *S*|_*l*_. Let *p, q*∈ℝ_≥1_. Let Inj(*l, k*) be the set of injections from *l* into *k*, that is the functions *σ* such that for all *i < l, σ*(*i*) ≤ *k*. We define the following function:

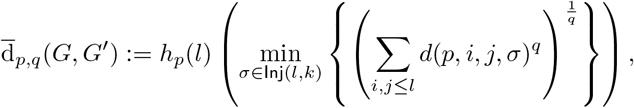

and similarly for 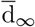. Here, *h*_*p*_(*l*) is a function of the number of species, *l*, in the smaller ecosystem, for a given *p*, that ensures *d*(*p, i, j, σ*) has the same dimensions for all numbers of living compartments.

Informally, Definition 3.5 finds the closest matching smaller matrix within the larger matrix for comparison.

## 4 Example 1: Jellyfish

In this section, we show how our approach can provide understanding of the differences in dynamics between models in a small test case. We compare four models for the dynamics of a system consisting of only two taxa: polyps of the moon jellyfish *Aurelia aurita* and an amalgamation of all their potential competitors. We show that our approach distinguishes between models in a way that is different from considering their fit to observed data.

### 4.1 Study system

Polyps of the moon jellyfish *Aurelia aurita* are believed to compete with other sessile marine organisms on hard substrates (Ishii and Katsukoshi, 2010). To learn more about the nature of this competitive interaction, the development of communities of *A. aurita*, other organisms such as solitary and colonial ascidians and bryozoans (amalgamated into ‘potential competitors’, which we treat as a homogeneous group) on PVC settlement panels was studied over an 8 week period (Boughton et al., 2023). Two treatments were applied: removal of half the *A. aurita* every week; and removal of half the potential competitors every week. As expected, removal of potential competitors increased the relative abundance of *A. aurita*. However, removal of *A. aurita* unexpectedly appeared to decrease the relative abundance of potential competitors. Experimental panels were placed at either 1 m or 3 m depth. For simplicity, we only consider the 1 m panels here.

### 4.2 Models

Four differential equation models were fitted in an attempt to understand the dynamics of the community. The original models included immigration from outside the system, but we include only internal dynamics here because our theoretical approach is based on closed systems (and we therefore exclude a fifth model, identical to the basic model described below except in immigration). The state variables for all models are *x*_1_ (the proportion of space occupied by potential competitors, denoted by *x* in the original paper) and *x*_2_ (the proportion of space occupied by *A. aurita* polyps: denoted by 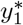 in the original paper), with *x*_1_ *>* 0, *x*_2_ *>* 0 and *x*_1_ + *x*_2_ ≤ 1. We write all the models in Kolmogorov form:

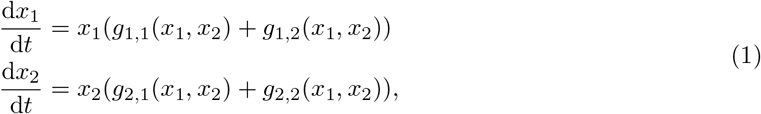

where *g*_*ij*_ is the effect of *j* on the proportional population growth rate of *i*. The models, with the form of each function in Equation 1, are given in Table 1. The overgrowth model appeared to fit the data better than the other three models, although none was able to fully reproduce the pattern of treatment effects (Boughton et al., 2023). Note that in some cases the choice of how to allocate terms in the model to the *g*_*ij*_ in Equation 1 was not obvious. For example, in the predator protection model, we put the term 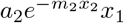, representing a reduction in predator mortality of potential competitors due to protection by polyp tentacles, into the self effect *g*_1,1_. We made this choice because the term is linear in *x*_1_ but nonlinear in *x*_2_ and represents a modification of mortality rather than a direct interaction. Similarly, in the growth facilitation model, we put the terms 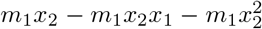, representing facilitation of growth of potential competitors onto unoccupied space by the presence of polyps, into the nonself effect *g*_1,2_. We made this choice because the term represents a modification by polyps of the mechanism of competition for space between groups of organisms.

**Table 1:** Models for interactions between *A. aurita* polyps *x*_2_ and potential competitors *x*_1_, with general form given by Equation 1. Parameters and dimensions: *a*_1_ (T^*−*1^), proportional rate at which proportion of unoccupied substrate is reduced by growth of potential competitors already on the substrate; *a*_2_ (T^*−*1^), proportional rate at which proportion of unoccupied substrate is increased by death of potential competitors already on the substrate; *b*_1_ (T^*−*1^), proportional rate of increase of polyp number on substrate by budding; *b*_2_ (T^*−*1^), proportional death rate of polyps on substrate; *m*_1_ (T^*−*1^): increase in rate of growth of potential competitors onto unoccupied space for a unit increase in proportion of space occupied by polyps; 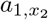 (numbers^*−*1^L^2^ T^*−*1^), rate at which potential competitors overgrow polyps; *m*_2_ (T^*−*1^, multiplied by a unit of time representing the time needed for a predator to travel along a given path), rate at which predators encounter polyps.

| Model | $g_{1,1}$ | $g_{1,2}$ | $g_{2,1}$ | $g_{2,2}$ |
| --- | --- | --- | --- | --- |
| basic | $a_1 + a_2 - a_1x_1$ | $-a_1x_2$ | $-b_1x_1$ | $b_1 + b_2 - b_1x_2$ |
| growth facilitation | $a_1 + a_2 - a_1x_1$ | $-a_1x_2 + m_1x_2 - m_1x_2x_1 - m_1x_2^2$ | $-b_1x_1$ | $b_1 + b_2 - b_1x_2$ |
| overgrowth | $a_1 + a_2 - a_1x_1$ | $-a_1x_2 + a_{1,x_2}x_2$ | $-b_1x_1 - a_{1,x_2}x_1$ | $b_1 + b_2 - b_1x_2$ |
| predator protection | $a_1 + a_2e^{-m_2x_2} - a_1x_1$ | $-a_1x_2$ | $-b_1x_1$ | $b_1 + b_2 - b_1x_2$ |

### 4.3 Parameter estimation, pseudodistance calculation and visualization

For each model, parameters were estimated using Bayesian methods, as described in Boughton et al. (2023). We took 1000 draws from the posterior distributions of parameters from each model. For each draw and each pair of models, we calculated the pseudodistances from Definition 3.3 as follows. The matrices whose elements are given in Table 1 are matrices of 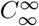 functions, so we can measure pseudodistances between them using 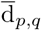 (though they need not correspond to ecosystem models in the sense of Section 6, as they may violate the axioms there required). Let *g*_*i,j*_ and 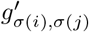 denote a pair of corresponding functions from Table 1, from models *M* and *M*′ respectively, with *i, j* ∈ *{x*_1_, *x*_2_*}* and *σ* ∈ Perm(2) being one of the two possible permutations of (*i, j*) (even though in this case we know that the set of species is the same in the two models). Then for each such pair of functions, we computed *d*(*p, i, j, σ*):

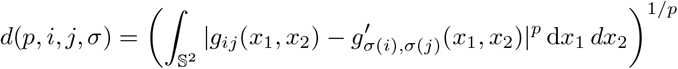

for *p* equal to 1 or 2, where S^2^ denotes the standard 2-simplex. We computed integrals over the simplex using adaptive integration in the R package `SimplicialCubature`version 1.3 (Nolan et al., 2021). We then summed over pairs of functions and minimized over permutations of matching species to obtain

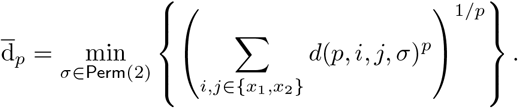

For *p* = ∞, we computed *d*(∞, *i, j, σ*):

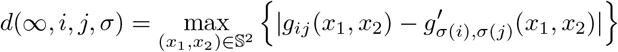

using the `constrOptim()` function in R (R Core Team, 2023), and then

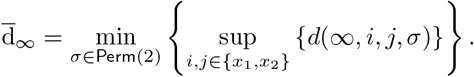

No normalising function was required in this case because the *x*_*i*_ are all dimensionless. Computing all pseudodistances in R version 4.1.2 (R Core Team, 2023) took approximately 130 min on a virtual server with 4 Intel Xeon Gold 6242R 3.10 GHz cores and 16 GiB RAM.

We visualized the distribution of pseudodistances among models using classical multidimensional scaling in two dimensions. In these data, each pseudodistance was minimized by the identity permutation, so each is in fact a distance. A distance matrix among *n* objects can be exactly represented in a Euclidean space if and only if the centred inner product matrix **B** = **HAH** is positive semidefinite, where **A** is a matrix of −1*/*2 times the squared distances, and **H** = **I** − *n*^−1^**11**′, where **I** is the *n × n* identity matrix, and **1** is a row vector of *n* 1s (Mardia et al., 1979, Theorem 14.2.1). This condition was met for 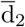 but not always for 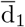 or 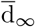. Nevertheless, the classical scaling algorithm is optimal in the sense of minimizing the squared discrepancy between original and fitted centred inner product matrices even if the centred inner product matrix is not positive semidefinite (Mardia et al., 1979, Theorem 14.4.2). We measured the goodness of fit of the two-dimensional representations using the quantity 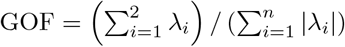 (Mardia et al., 1979, p. 408), where *λ*_*i*_, *i* = 1, 2, …, *n* are the eigenvalues of **B**. We plotted the two-dimensional representations for each draw from the posterior for each model, and calculated the value of GOF, using the function `cmdscale()` in R (R Core Team, 2023). All pseudodistances produced similar patterns. We therefore report results for 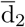 in the main text, and 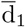 and 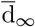 in appendices.

### 4.4 Differences between models

For each of the three pseudodistances, the overgrowth model (Figures 3, A1 and A2, purple; Tables 2, A1 and A2, third row and column) had relatively distinct dynamics from the other models. This is consistent with the original analysis (Boughton et al., 2023), in which the overgrowth model appeared to be the best fit to the data (measured in terms of expected log predictive density across all experimental panels, including those at 3 m, not considered here). However in the original analysis the other models were almost indistinguishable, but our new analysis gave some evidence of a systematic difference in dynamics between the basic (Figures 3, A1 and A2, green) and predator protection (Figures 3, A1 and A2, pink) models. Comparisons in terms of fit to data and dynamics can give different results because they evaluate different aspects of model behaviour. When evaluating fit to data, all models were initialized with an empty habitat, corresponding to the initial condition in the experiment. Furthermore, both observed and predicted dynamics only explored a small part of the simplex. In contrast, the comparisons of dynamics reported here integrate over the whole of the simplex, and thus give a fuller picture of differences in model behaviour.

**Table 2:** Pseudodistances 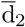 between 4 models for a sessile community (basic, growth facilitation, over-growth, predator protection). Posterior means from 1000 draws, with 0.025 −and 0.975 −quantiles in brackets.

|  | basic | growth facilitation | overgrowth | predator protection |
| --- | --- | --- | --- | --- |
| basic | 0.00 (0.00, 0.00) | 0.05 (0.02, 0.09) | 0.14 (0.07, 0.23) | 0.05 (0.02, 0.09) |
| growth facilitation | 0.05 (0.02, 0.09) | 0.00 (0.00, 0.00) | 0.14 (0.06, 0.22) | 0.05 (0.02, 0.10) |
| overgrowth | 0.14 (0.07, 0.23) | 0.14 (0.06, 0.22) | 0.00 (0.00, 0.00) | 0.15 (0.07, 0.23) |
| predator protection | 0.05 (0.02, 0.09) | 0.05 (0.02, 0.10) | 0.15 (0.07, 0.23) | 0.00 (0.00, 0.00) |

**Figure 3:**
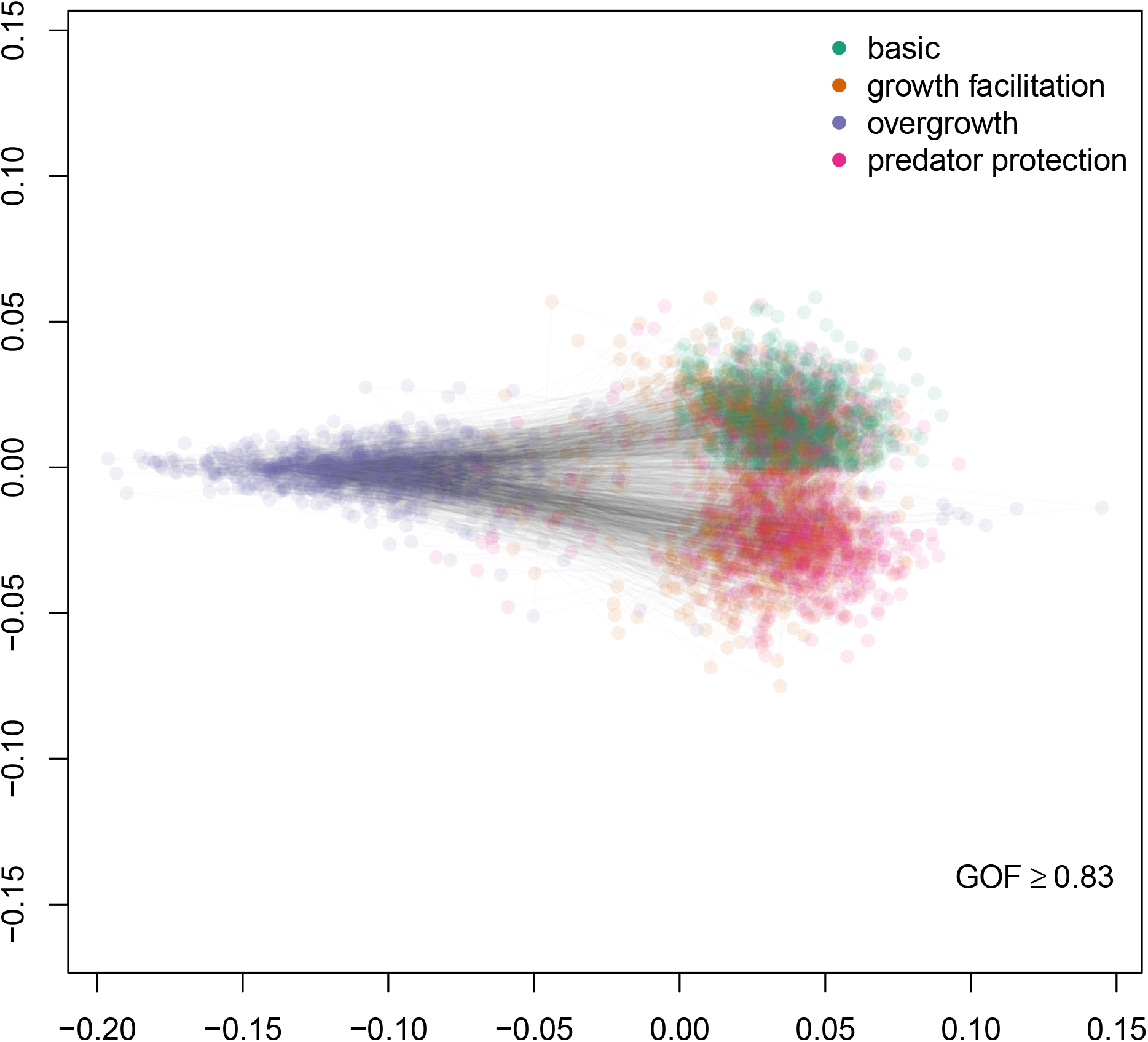
Classical multidimensional scaling of pseudodistances 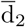 between 4 models for a sessile community. Each set of 4 coloured points (green: basic, orange: growth facilitation, purple: overgrowth, pink: predator protection) connected by lines represents a single pseudodistance matrix with parameters drawn from their posterior distributions (1000 draws in total). Goodness-of-fit (GOF) for the two-dimensional representation was at least 0.83 in all draws.

## 5 Example 2: ecosystem models from the enaR troModels database

In this section, we show how our approach can in principle be used to study patterns in the dynamics of large ecosystem models. We calculate pseudodistances between members of a set of 50 freshwater and marine ecosystem models containing 4 to 122 living compartments (Borrett and Lau, 2014). We make some major assumptions to get from static energy flow networks to models for dynamics, and our results should therefore be taken as proof of principle, rather than necessarily being realistic. We show that freshwater and marine ecosystem models appear to have somewhat distinct dynamics, although the freshwater models may represent only a few different ecosystems. We also show that different versions of the same ecosystem model (e.g. from different sites or times) may have very substantially different dynamics.

### 5.1 Ecosystem dynamics from flow networks

Let *x*_*i*_ be the stock of energy in the *i*th compartment of a flow network, in which each compartment is either non-living or living. Let *f*_*ij*_ be the flow of energy from compartment *j* to compartment *i, z*_*i*_ be the rate of external input to compartment *i, e*_*i*_ be the rate of export from compartment *i*, and *r*_*i*_ be the rate of energy dissipation from compartment *i*. These are the quantities given in a typical ecosystem flow network (e.g. Borrett and Lau, 2014). It is usual to work with balanced networks, in which each stock is constant over time. For example, the balanced flow network for the freshwater Cone Spring ecosystem (Tilly, 1968) is shown in Figure 4.

**Figure 4:**
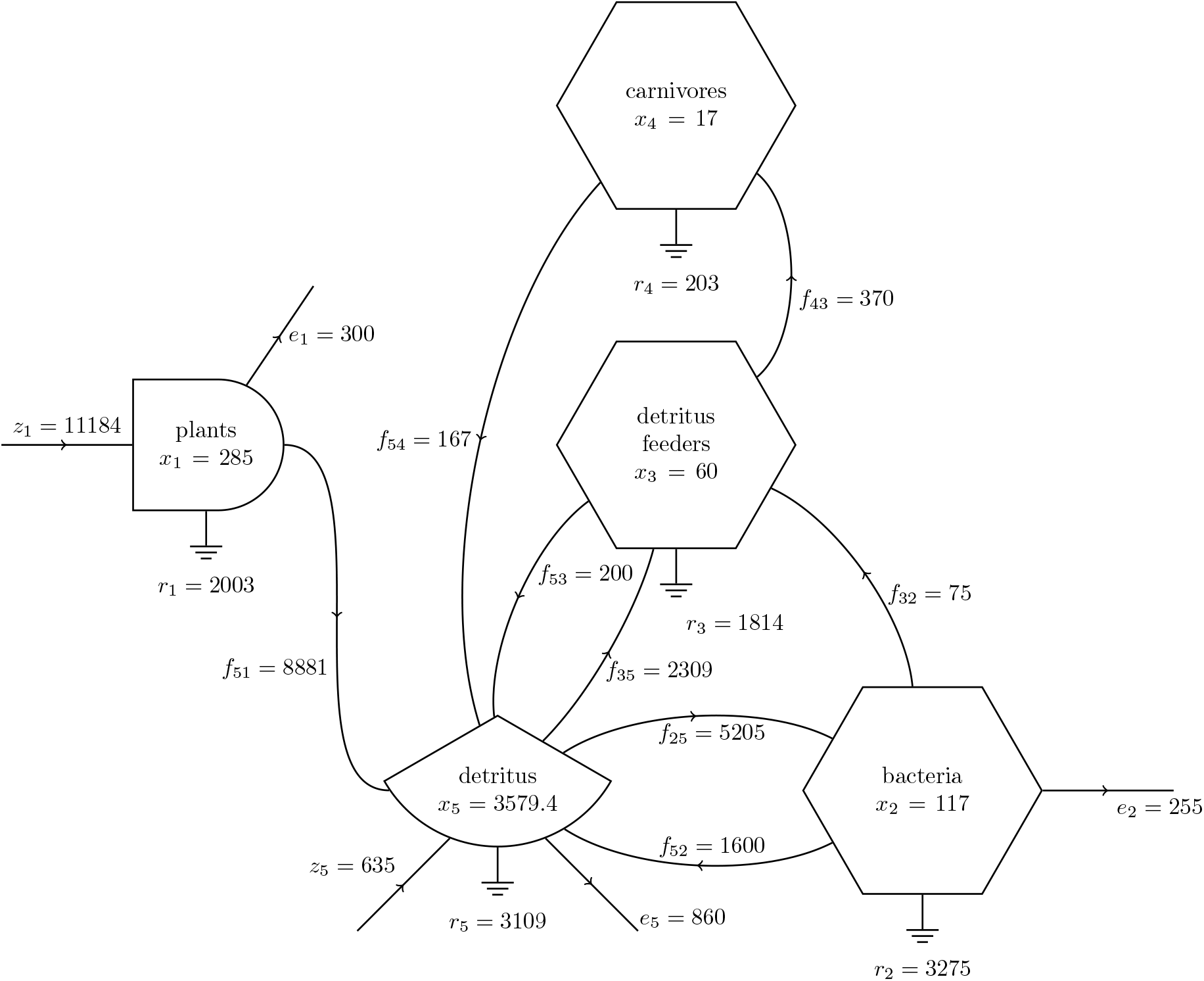
Flow network for the Cone Spring ecosystem. Data from the compilation by Borrett and Lau (2014), originally collected by Tilly (1968). Symbols are based on Odum (1983, p. 8): bullets represent producers, hexagons represent consumers, circle segments represent nonliving tanks, arrows represent flows of matter or energy, ground symbols represent respiration. In each compartment, *x*_*i*_ represents the stock in kcal m^−2^. Edge weights are in kcal m^−2^ a^−1^: *f*_*ij*_ represents a flow from compartment *j* to compartment *i, z*_*i*_ represents a flow into compartment *i* from outside the system, *e*_*i*_ represents a flow out of the system from compartment *i*, and *r*_*i*_ represents the rate of energy dissipation by compartment *i*.

We want to compare the ecosystem dynamics generating flow networks, in contrast to analyses treating such networks as static objects whose structure is of interest (e.g. Patten, 1978; Ulanowicz, 1986). It has been claimed that most aspects of ecosystem flow networks can be described by linear models in which flows are proportional to upstream stocks (Odum, 1983, p. 11). This claim does not appear to have a strong justification (Månsson and McGlade, 1993). Instead, we assume that balanced ecosystem flow networks are a phenomenological description of energy flows at equilibrium, and choose simple but plausible differential equation models for the underlying mechanisms. The energy flow diagrams themselves do not contain enough information to uniquely determine the differential equations, so our results should be viewed only as proof of concept. Following a similar approach to DeAngelis (1980, eq. 20), we assume that the dynamics of the *i*th stock are given by the differential equation

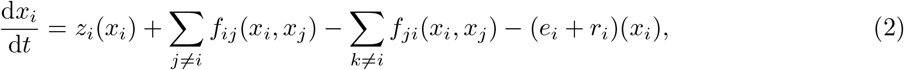

where the inputs *z*_*i*_, outputs *e*_*i*_ and dissipation rates *r*_*i*_ are functions of the stock *x*_*i*_, and the rate *f*_*ij*_(*x*_*i*_, *x*_*j*_) of energy flow from the *j*th compartment to the *i*th compartment are functions of the stocks in compartments *i* and *j*. For non-living destination compartments, we assume that the flow is proportional to the source stock (or is a constant for external inputs), as in Odum (1983, p. 11). However, for living destination compartments, we assume a mass-action model in which the flow is proportional to the product of source and destination stocks (or to the destination stock for external inputs, or to the source stock for exports and dissipation). This is a Lotka-Volterra model for interactions between consumers and resources, and is likely to be a good approximation more often than dependence on the source stock alone, although such “donor-controlled” interactions do occur for some living destination compartments (Pimm, 1982, section 5.2). For external inputs to living compartments, assuming that the flow is proportional to the destination stock is appropriate if these inputs represent energy fixation (e.g. by photosynthesis) rather than transport of organisms into the system. Other approaches to conversion of flow networks into differential equations are possible (e.g. Walters et al., 1997), but generally require additional information beyond the flow network alone. The form of model for each type of flow is summarized in Table 3. We calculate the parameters based on stocks and flows in the balanced network. For example, given *f*_*ij*_, *x*_*i*_ and *x*_*j*_ in the balanced network, *α*_*ij*_ = *f*_*ij*_*/*(*x*_*i*_*x*_*j*_).

**Table 3:** Models for energy flows between compartments in an ecosystem flow network. Here, *x*_*i*_ represents the stock in compartment *i* (dimensions ML^−2^ of carbon), *z*_*i*_(*x*_*i*_) represents an external input into compartment *i, e*_*i*_(*x*_*i*_) represents an export from compartment *i, r*_*i*_(*x*_*i*_) represents a dissipation from compartment *i*, and *f*_*ij*_(*x*_*i*_, *x*_*j*_) represents a flow from *j* to *i*. All flows *z*_*i*_, *e*_*i*_, *r*_*i*_, *f*_*ij*_ have dimensions ML^−2^T^−1^ of carbon. The parameters *ζ*_*i*_, *ℰ*_*i*_, *ρ*_*i*_ and *α*_*ij*_ are determined by the stocks and flows, under the assumptions explained in the text.

| Flow type | Source compartment | Destination compartment | Form |
| --- | --- | --- | --- |
| External input $z_i(x_i)$ to $i$ | - | non-living | $z_i(x_i) = \zeta_i$ |
| | - | living | $z_i(x_i) = \zeta_i x_i$ |
| Export $e_i(x_i)$ from $i$ | non-living | - | $e_i(x_i) = \varepsilon_i x_i$ |
| | living | - | $e_i(x_i) = \varepsilon_i x_i$ |
| Dissipation $r_i(x_i)$ from $i$ | non-living | - | $r_i(x_i) = \rho_i x_i$ |
| | living | - | $r_i(x_i) = \rho_i x_i$ |
| Flow $f_{ij}(x_i, x_j)$ from $j$ to $i$ | non-living | non-living | $f_{ij}(x_i, x_j) = \alpha_{ij} x_j$ |
| | living | non-living | $f_{ij}(x_i, x_j) = \alpha_{ij} x_j$ |
| | non-living | living | $f_{ij}(x_i, x_j) = \alpha_{ij} x_i x_j$ |
| | living | living | $f_{ij}(x_i, x_j) = \alpha_{ij} x_i x_j$ |

When comparing ecosystem flow networks, we restrict our attention to living compartments. If *i* is a living compartment, then under the assumptions about flow networks from Table 3, Equation 2 is of the Kolmogorov form

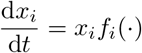

for some function *f*_*i*_, and we partition the pairwise effects as follows:

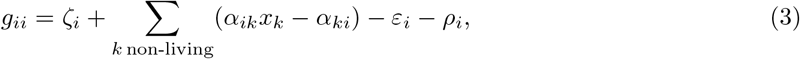

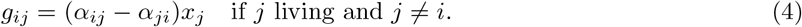

For flows between living compartments *i* and *j, g*_*ij*_ is the net proportional effect of *j* on *i*. In contrast, if *i* is a non-living compartment, then under the assumptions from Table 3, Equation 2 will not be of Kolmogorov form, because flows into *i* do not depend on the stock *x*_*i*_, and it is not obvious how to choose an appropriate *g*_*ii*_. Since flows to and from non-living compartments affect the dynamics of a living compartment *i*, they must therefore be included in the self effect *g*_*ii*_. To make the *g*_*ii*_ functions of *x*_*i*_ alone, we fix *x*_*k*_ for each *k* non-living in Equation 3 at its equilibrium value in the calculations below.

For the living compartments in the Cone Spring ecosystem (Figure 4), we have

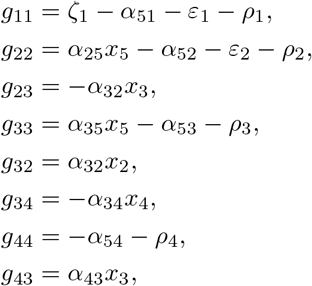

where parameters are calculated using Table 3, parameters not listed are zero, and functions not listed are zero functions.

### 5.2 Pseudodistances

As in Section 4, the matrices whose elements are given in Equations 3 and 4 are matrices of 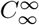 functions, so we can measure pseudodistances between them using 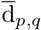. Also as in Section 4, the matrices need not correspond to ecosystems models in the sense of Section 6, as they may violate the axioms there required. For a pair of ecosystem models, let *V* be the set of living compartments in the ecosystem model with fewer living compartments, *W* the set of living compartments in the ecosystem model with more living compartments, and Inj(*V, W*) be the set of injective mappings of living compartments of *V* to living compartments of *W*. Let *g*_*ij*_ and 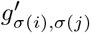 denote corresponding pairwise effect functions in the two ecosystems, for *σ* ∈ Inj(*V, W*). Then the distance between these two functions is given by

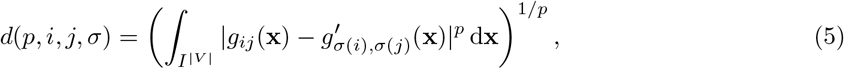

where *I* = [0, *κ*], *κ* is the largest stock observed in any ecosystem. The choice of *κ* is somewhat arbitrary, but some value is needed for the integral to be defined. To compare between ecosystems we need the same value in all cases, and the largest observed stock across all ecosystems is the obvious choice. We integrate over the whole of **x** even though in this particular class of models, *g*_*ij*_ is a function of *x*_*j*_ alone, and *g*_*ii*_ is not a function of either *x*_*i*_ or *x*_*j*_, for consistency with our general definition.

Then if *i* and *j* are both living, *i ≠ j* and *p ≠* ∞,

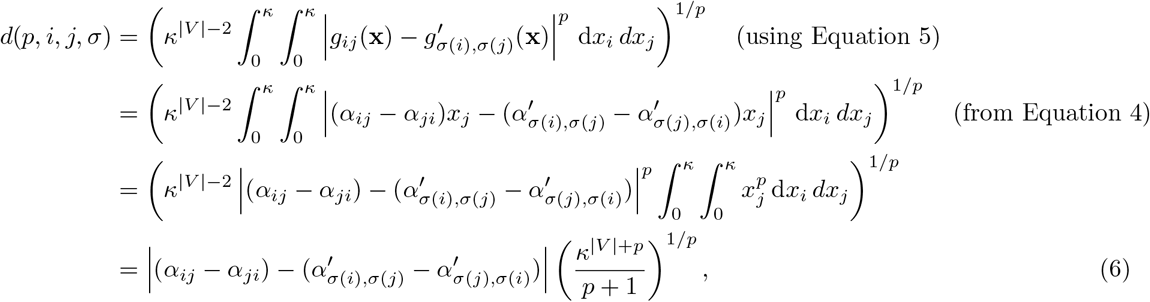

where *α*_*ij*_ and 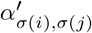 denote corresponding parameters in the two ecosystems. Letting *p*→ ∞ in Equation 6,

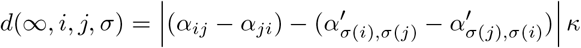

if *i* and *j* are both living and *i ≠ j*.

Similarly, if *i* is living, *i* = *j* and *p ≠* ∞,

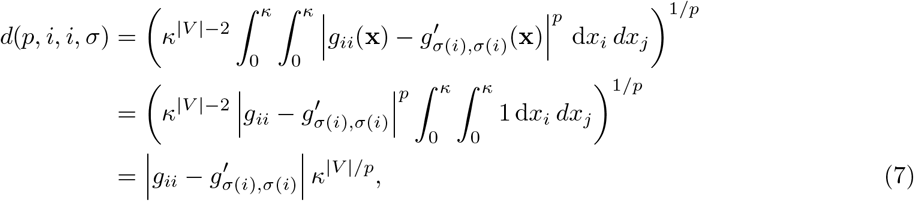

where *g*_*ii*_ and 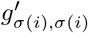 denote corresponding self effects in the two ecosystems, given in Equation 3. Letting *p* → ∞ in Equation 7,

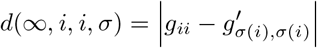

for *i* living and *i* = *j*.

We set *q* = *p* in Definition 3.2. Thus for *p ≠* ∞,

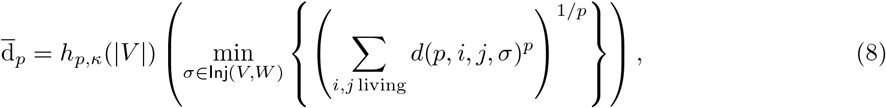

where *h*_*p,κ*_(|*V* |) is a normalising function of the number of living compartments in the smaller ecosystem that ensures 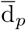 has the same dimensions for all numbers of living compartments. Here, each *x*_*i*_ has dimensions ML^−2^ of carbon (Table 3). A natural choice for *h*_*p,κ*_(|*V* |) is therefore

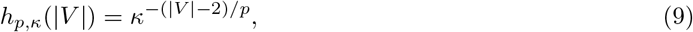

so that *d*(*p, i, j, σ*) will have dimensions M^2*/*p^L^−4*/*p^T^−1^ of carbon, for any |*V* |.

For *p* = ∞,

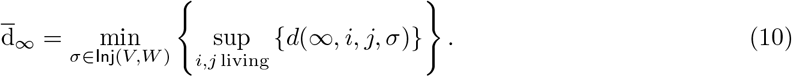

We can set *h*_*p,κ*_(|*V* |) = 1 in Equation 10 because the dimensions do not depend on |*V* |.

Note that for the models considered here, *d*(*p, i, j, σ*) = *d*(*p, j, i, σ*) because of the symmetry in Equation 6, but this is not true for all ecosystem models, so we include both in the summation in Equation 8, as in the general Definition 3.2.

### 5.3 Data sources

The `troModels` data set in the R package `enaR` version 3.0.4 (Borrett and Lau, 2014) contains 59 ecosystem models ordered by increasing number of compartments. We included only living compartments, defined by the `living` variable for each model in `tromodels`, as justified in section 5.1. Stocks and flows in these models are given in a variety of units. We converted all flows to mgm^−2^d^−1^ of carbon, and all stocks to mgm^−2^ of carbon, using conversion factors from Odum (1971, p. 39, Table 3-1, for ash free dry weight to kilocalories) and Peters (1983, Appendix Ia, for all other conversions). Odum (1971, p. 39, Table 3-1) lists kilocalories per gram ash free dry weight for terrestrial plants, algae, invertebrates excluding insects, insects and vertebrates: we used the mean of these values for all such conversions. Peters (1983, Appendix Ia) gives a range of 3 kg to 10 kg wet mass per kg dry mass: we used the midpoint of this range for all such conversions. For the Mdloti estuary model (ecosystem 40), the `enaModelInfo` object gave the units of flow as kg capita^−1^ a^−1^ of carbon, while the original source (Scharler, 2012) and the package vignette (Lau et al., 2023) both gave the units as mgm^−2^d^−1^ of carbon. We assumed that the original source was correct (and version 3.1.1 of enaR now agrees with the source). For any models that were not already balanced, we applied the `balance()` function from `enaR` with the `AVG2` algorithm (Allesina and Bondavalli, 2003). We removed any compartments with zero stock. In principle, such compartments could be included, but our assumptions about the dependence of stocks on flows (Table 3) would lead to divide-by-zero errors. We also excluded 9 ecosystem models (numbers 1, 2, 3, 4, 5, 7, 8, 13 and 23) for which all stocks were given as 1 unit in the original units, which is unlikely to be correct. In many of these, such as the lake ecosystem models 2, 3, 4 and 5 (Richey et al., 1978) the stocks were not given in the original sources, while in others such as the English Channel model (ecosystem 8, Brylinsky, 1972) stocks were given in the original source and were not all 1 unit. This left 50 ecosystem models, listed in Table A3, with between 4 and 122 living compartments (median 29.5). We set the upper limit of integration *κ* in distance calculations to the largest living stock in any included ecosystem, which was 1.07 × 10^7^ mgm^−2^.

We classified each ecosystem as predominantly either marine (40 models) or freshwater (10 models) using information in the original sources wherever possible (the only exceptions were models 50 and 51 for cypress wetlands in Florida, and 56 and 57 for mangroves in Florida). We classified any ecosystem with clear marine influence, including tidal marshes, intertidal habitats, estuaries, brackish bays and mangrove ecosystems, as marine. We classified all other ecosystems as freshwater, although some, such as cypress and graminoid wetlands in Florida, included terrestrial components (Heymans et al., 2002).

### 5.4 Computation

Let *V* be the set of living compartments in the ecosystem model with fewer living compartments, *W* the set of living compartments in the ecosystem model with more living compartments, and Inj(*V, W*) be the set of injective mappings of living compartments of *V* to living compartments of *W*. As in Definition 3.5, we want to compute min_Inj(*V,W*)_ 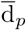. This is a version of the quadratic assignment problem, which is NP-hard (Burkard et al., 1998). Of the |*W*|! permutations of *W*, only the order of the first |*V* | elements matters, so | Inj(*V, W*)| = |*W*|!*/*(|*W*|−|*V* |)!. For | Inj(*V, W*)| ≤ 3.41*×*10^5^, we enumerated all mappings and selected the one giving the smallest 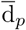. For pairs of models with larger numbers of possible mappings, we used simulated annealing implemented in the R function `optim()` with the argument `method = “SANN”`, with 10 random start points and arguments `maxit = 3e4, temp = 10, tmax = 20.` At each iteration, a new candidate mapping was generated by permuting the indices of compartments in the larger ecosystem, and using only the first |*V* |of these. We chose to switch from enumeration to simulated annealing at 3.41 *×* 10^5^ mappings by running both enumeration and simulated annealing on a small set of 5 ecosystems with between 120 and 3628800 mappings, using least-squares regression to estimate the relationship between log time and log number of mappings for each method, and calculating the number of mappings at which the predicted time was less for simulated annealing than for enumeration. We checked that simulated annealing found the same solutions as enumeration in these small cases. With the argument `maxit=3e4`, there was generally no change in the best solution found for pairs of small ecosystems after 1000 iterations, but improvements were sometimes seen after 2000 iterations in pairs of larger ecosystems. Computing all pseudodistances in `R` version 4.1.2 (R Core Team, 2023) took approximately 1248 hours on a virtual server with 4 Intel Xeon Gold 6242R 3.10 GHz cores and 16 GiB RAM.

### 5.5 Visualization

We visualized patterns in pseudodistances using non-metric multidimensional scaling (Kruskal, 1964), which is suitable for pseudometrics such as 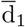, 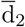 and 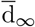 because it uses only rank information. We used the implementation in the function `metaMDS()` in the `R` package `vegan` version 2.6-2 (Oksanen et al., 2022), with default options, including the underlying `monoMDS` engine, which is able to handle zero pseudodistances. We plotted stress against dimension, and ordinations in 2 and 3 dimensions. We distinguished marine and freshwater ecosystems on the ordinations using colours. Where a single source reported multiple models for the same ecosystem at different times and/or sites, we drew lines connecting the points representing these models.

### 5.6 Patterns of dissimilarity

In general, two-dimensional non-metric multidimensional scaling (Figure 5) gave poor representations of the pseudodistances for 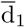, 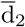 and 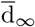, in terms of the rule of thumb for interpreting stress from Kruskal (1964, p. 3). However, increasing the number of dimensions did not give a dramatic improvement (Figure A3), suggesting that these data were fundamentally high-dimensional. Three-dimensional non-metric multidimensional scaling (Figure A4) was harder to visualize, but contained essentially the same patterns. These patterns also did not differ substantially between 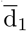, 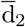 and 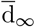 (Figures 5 and A4, panels). It is important not to overinterpret the patterns in these ordinations given the high stress. Nevertheless, we comment on some features.

**Figure 5:**
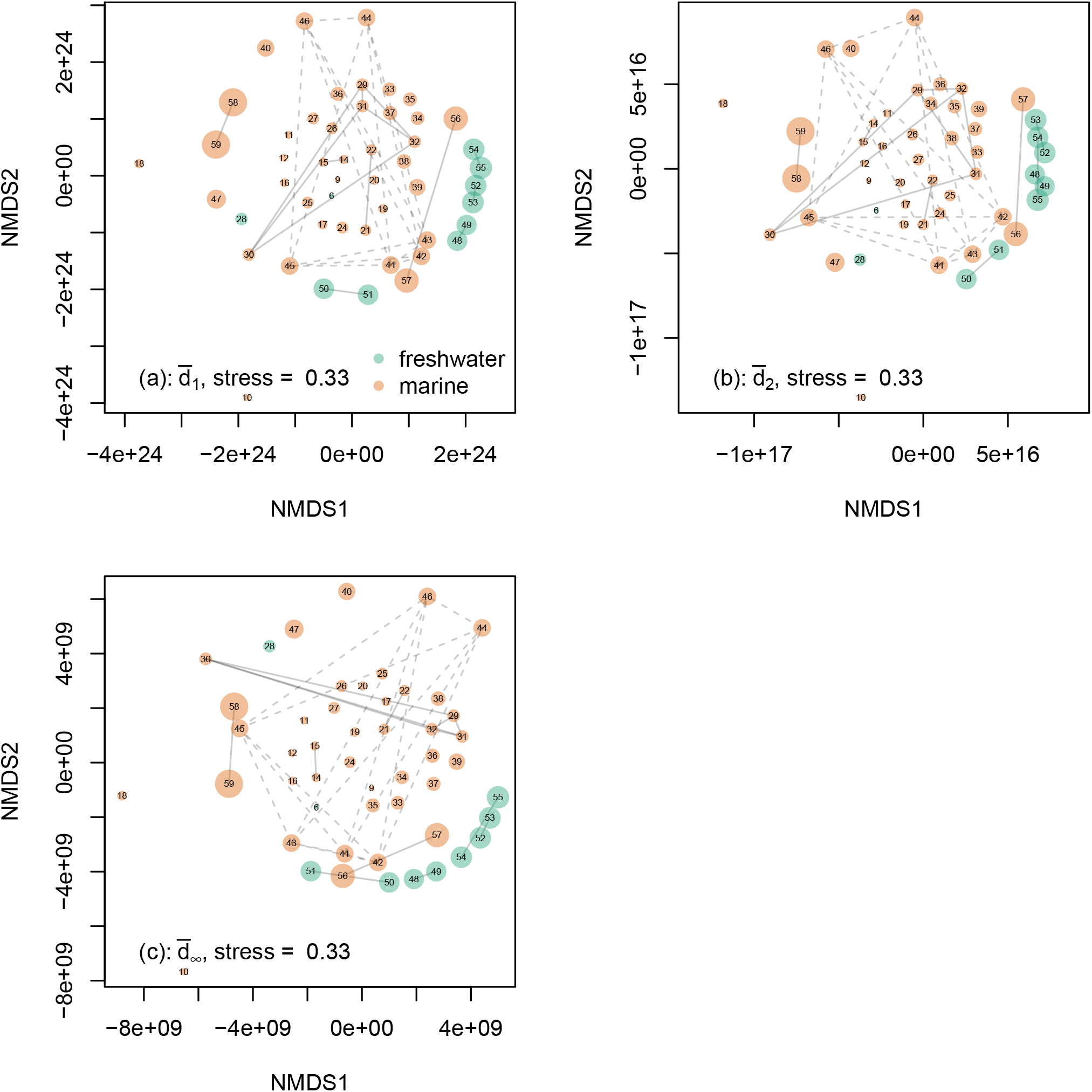
Two-dimensional non-metric multidimensional scaling of pseudodistances (a) 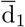, (b) 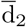 and (c) 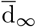 among 50 ecosystem models from the troModels compilation in the enaR package (Borrett and Lau, 2014). Numbers on the ordinations correspond to model numbers in troModels. The stress is indicated for each ordination. Point areas are proportional to the number of living compartments in the model. Point colours represent habitat types (green freshwater, orange marine). Lines connect models for different versions of the same ecosystem such as sites or times. Dashed lines indicate the large cluster of St Marks Seagrass ecosystems at different sites and times, numbers 41 to 46 (Baird et al., 1998).

The most obvious pattern in both ordinations was that ecosystems with larger numbers of living compartments (Figures 5 and A4, larger points) tended to be far apart from each other around the edges of the cloud of points. The pseudodistances 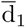 and 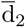, which involve summation of functional metrics between pairwise effects over all corresponding living compartments in a pair of ecosystems (Equation 8). For two ecosystems with larger numbers of living compartments, the summation will be over a large number of functional metrics, and thus a larger pseudodistance is expected. The pseudodistance 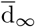 is just the maximum of these functional metrics over pairs of living compartments (Equation 10), and would be expected to be larger if taken over more pairs, other things being equal.

Most of the freshwater ecosystems (Figures 5 and A4, green points) tended to form a cluster separate from the marine ecosystems (Figures 5 and A4, orange points). However, the freshwater cluster contained relatively few distinct ecosystems: Florida graminoid marshes in the wet and dry seasons (ecosystems 48 and 49); the nearby Florida cypress swamps in the wet and dry seasons (ecosystems 50 and 51), which have many species in common with the graminoid marshes (Heymans et al., 2002); and two trophically similar lake ecosystems (Miehls et al., 2009a), Lake Oneida (ecosystems 52 and 53) and Bay St Quinte, Lake Ontario (ecosystems 53 and 54) before and after zebra mussel invasion. The two remaining freshwater ecosystems, Cone Spring (Tilly, 1968) and Okefenokee Swamp (Whipple and Patten, 1993) did not form part of this freshwater cluster, and it remains to be seen whether the cluster is a genuine pattern.

Different versions of the same ecosystem (e.g. different sites or times: connected by lines in Figures 5 and A4) were sometimes but not always similar. For example, the pairs in the freshwater cluster discussed above, and the wet and dry season models for the Florida Bay lagoon ecosystem (ecosystems 58 and 59, Ulanowicz et al., 1998), were relatively similar. In contrast, in the set of Neuse Estuary models for early and late summer in the years 1997 and 1998 (ecosystems 29, 30, 31, and 32, Baird et al., 2004b), ecosystem 30 (late summer 1997) appeared very different from the the other models. The Neuse Estuary experienced severe hypoxia in the summer of 1997, with substantial changes in trophic structure (Baird et al., 2004b). In the set of 6 models for the St Marks Seagrass ecosystem (ecosystems 41 to 46, Baird et al., 1998), models 45 and 46 correspond to a pair of sites subject to stronger currents and more variable salinity than the other models, while models 41, 43 and 45 are from January 1994 and models 42, 44 and 46 are from February 1994, with higher water temperatures and immigration of fish and birds (Baird et al., 1998). There is substantial variation between models (Figures 5 and A4, dashed lines), but they did not segregate simply by date or by site group alone.

## 6 Spaces and Natural Measurements of Ecosystems

Here, we outline an axiomatic approach to community ecology, by specifying a class of models of ecosystems, the *limitative ecosystems*, or *L-ecosystems* (Section 6.1). In Section 6.2, we define pseudometrics on these ecosystems. Our main result, in Section 6.4, is that under a natural notion of a “type” of ecosystem, based on a natural measurement, there is only one type of ecosystem. Appendix E contains the proofs.

### 6.1 Limitative Axioms

For *L*-ecosystems, we specify a space of matrices of functions, which we call the *space of L-ecosystems*. We specify a set of matrices of functions, then state some axioms, which a matrix must satisfy to be an *L*-ecosystem. We motivate these axioms biologically.

Our limitative approach for axioms for models of ecosystems resembles e.g. Ansmann and Bollenbach (2021) and Hutchinson (1978, pages 1-5), following Lotka (1956, pages 64-65). Throughout, we assume that our system is biotically closed, i.e. that there is no emigration/immigration/recolonisation. We do not assume that our systems is abiotically closed. We allow a non-producer to not predate on any species in the ecosystem (e.g. its prey has died out). We allow producers to predate on other species (e.g. predators with symbiotic algae). In Appendix C, we show that the opposite approach, where we build up from small pieces, rather than biologically restricting the kinds of matrices of functions, yields the same set of ecosystems. Hence, these can also be made into topology-like spaces.

Let *k*∈ ℕ. An *L*-ecosystem, *e*, with species labelled by 1, …, *k*, consists of four pieces of data, *G*^*e*^, **x**^*e*^, *P* ^*e*^, and *c*^*e*^, which we define like so:

1. *G*^*e*^ is a *k*-by-*k* matrix of infinitely differentiable real-valued functions on ℝ^2^, representing the pairwise (proportional, Axiom I.) effects of one species on another, as a value of their abundances,
2. **x**^*e*^ is a *k*-length vector of non-negative real-valued functions, representing species abundances over time,
3. *P* ^*e*^ is a *k*-by-*k* matrix such that:
  a. the diagonal entries specify whether the species is a producer (either Prod or 0),
  b. the off-diagonal entries specify whether one species predates on another (either 1 or 0), and
4. *c*^*e*^ is a *k*-by-*k* matrix of infinitely differentiable non-negative real-valued functions of ℝ_≥0_, specifying the efficiency of the conversion of units of one species into another in the case of predation.

Firstly, we write down the set of all such collections of data. Then we state the limitative axioms, with biological explanations. Let *k* ∈ ℕ. We call *e* a *candidate model for k* if 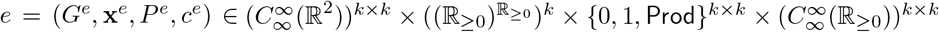. If *e* is clear from context, we drop the superscript *e*. Note that 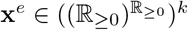 as it is a *k*-length vector of abundances with respect to time, so is a function non-negative reals to non-negative reals.

#### Definition 6.1.

Let *k* ∈ ℕ, and *e* be a candidate model for *k*. We call *e* an *L-ecosystem (over k)* if for all *i, j* ≤ *k*, and for all *x, y, t* ∈ ℝ_≥0_, the following hold.

A. (Postulate of Parenthood) For all *ϵ >* 0, if *x*_*i*_(*t*) = 0 then *x*_*i*_(*t* + *ϵ*) = 0. This means that if a species goes extinct, then it cannot come back from extinction at a later date, e.g. there is no recolonisation.
B. (Prey Consumption and Predator Growth) If *P*_*i,j*_ = 1, then *g*_*i,j*_(*x, y*) *>* 0 *> g*_*j,i*_(*x, y*). This means that if species *j* predates on species *i*, then the effect of prey consumption on predator abundance is positive.
C. (Only Predation and Production) If *P*_*i,j*_ = 0 and *i ≠ j, g*_*i,j*_(*x, y*) = 0. This means that if a species does not predate on any other, then it does not directly affect the abundance of that other species, i.e. the only ecological interaction mechanism is predation.
D. (Monotonicity) If *P*_*i,j*_ = 1 then *g*_*i,j*_(*x, y*) is strictly increasing in *y* and *g*_*j,i*_(*x, y*) is strictly decreasing in *x*. This means that if one species predates on another, then the effect of prey consumption on predator abundance is increasing.
E. (Producers can Increase when Everything is Rare) If *P*_*i,j*_ = Prod then *i* = *j* and for all *l* ∈ *I*, there exist abundances *b*_*l*_ ∈ ℝ_≥0_, such that if *x*_*l*_(*t*) *< b*_*l*_, then *g*_*i*_(*x*_*i*_(*t*), *x*_*i*_(*t*)) *>* 0. This means that for a given producer, if each species is lower than a species-dependent level, that producer may increase in abundance.
F. (Non-producers need Prey) If *P*_*i,i*_ *≠* Prod then *g*_*i,i*_(*x, x*) *<* 0. This means that non-producers cannot, alone, increase their abundance.
G. (Producers are Self-Limiting) If *P*_*i,i*_ = Prod then *g*_*i,i*_(*x, x*) is strictly decreasing in *x*. This means that as the abundance of a producer population increases, its proportional rate of increase decreases.
H. (Inefficient Conversion) If *P*_*i,j*_ ≠1 then *c*_*i,j*_ = 0, otherwise *P*_*i,j*_ = 1 and 0 ≤ *c*_*i,j*_(*x*) *< x*. This means that predation is never 100% efficient.
I. (Rates of Change)

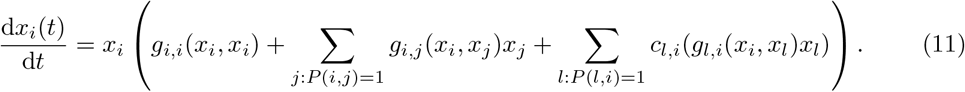

This means that 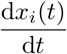 is governed by a Kolmogorov equation (see Kolmogorov, 1936; Sigmund,2007).

We denote the set of all *L*-ecosystems over *k* by *E*_*k*_. We define the set of *L*-ecosystems like so:

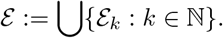

We call A. to I. the (limitative) axioms. As 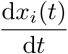 is a finite sum of continuous integrable functions, it is continuous and integrable; hence *x*_*i*_(*t*) is also continuous. Note that the pairwise proportional effect, *g*_*i,j*_, may be negative, but the solutions (in Axiom I.) are necessarily positive. These axioms allow for the species to evolve (*g*_*i,j*_ and *c*_*i,j*_ are not fixed), but forbid speciation (the number of species is fixed), and forbid changes in the predator-prey relationship (*P* (*i, j*) is fixed). Suitable variations of our set-up would permit such changes.

### 6.2 Pseudometrics on Models of Ecosystems

A model of an ecosystem consists of the data specified in Section 6.1. We now define a pseudometric on the set of ecosystems as in Section 3.2. We focus on the *dynamical* differences between ecosystem models, i.e. the matrix of pairwise proportional effects of species, so suppress the other three pieces of data. A simple elaboration of the resulting pseudometric would incorporate the other data. As it is no more difficult, we define the pseudometric on the set of candidate models.

#### Definition 6.2.

Fix a *k*∈ ℕ. Suppose that *e* and *f* are candidate models for *k, p* = ∞ or *p* ∈ ℝ_≥1_, and likewise for *q*. Let 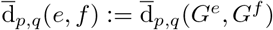.

There is a corresponding version of Definition 3.5 for candidate models with different numbers of species, again using *G*^*e*^ and *G*^*f*^ (as in Definition 6.2). Hence, we can form a topology-like family of sets on *E*:

#### Theorem 6.3.

Let *p, q* ∈ ℝ_≥1_. Then 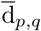 is a symmetric, non-negative function on *ℰ*^2^ such that 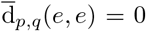 for all *e* ∈ *ℰ*. Moreover, 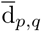 fails the triangle inequality, so is not a pseudometric on *E*. Let *τ*_*E,p,q*_ be the set of unions of open balls, 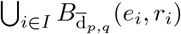, along with the empty set, ∅. Then *τ*_*E,p,q*_ is closed under unions, but need not be a topology.

If the functions *d* : *X*^2^ → ℝ_≥0_ from Theorem 6.3 are metrics, then this is a standard construction of a topology. However, in our case, they need not be metrics, but they are so-called *premetrics* (unhelpfully sometimes called pseudometrics). As a result, 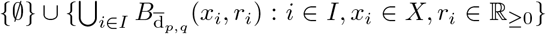 is not a topology, but it is closed under unions and contains ∅.

An alternative definition with premetrics, a *A* ⊆ *ℰ* is open if and only if for all *x ∈ A* there is an *r*_*x*_ *>* 0 such that 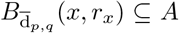 (see Arkhangel’skil and Fedorchuk, 1990, p. 23), is a topology on *ℰ*, but is essentially trivial: the balls 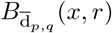 need not be open (this is because ecosystems may have different number of species, so the premetric strongly violates the triangle inequality).

### 6.3 Natural Measurements

An obvious question is what kinds of functions are continuous with respect to the family *τ*_*ℰ*_. In particular, we are interested in continuous functions *f* : *ℰ →* ℝ, which we call *natural measurements* on the set of ecosystems. Our family, *τ*_*ℰ*_, can only distinguish points based on their matrix of pairwise effects, *G*^*e*^ (i.e. if 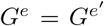 then ecosystems *e* and *e*′ are in exactly the same open balls, making *τ*_*ℰ*_ *non-Hausdorff*). Hence, a necessary condition for *f* : *ℰ*→ℝ_*≥*0_ to be continuous is that whenever 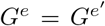, we have *f* (*e*) = *f* (*e*′).

Metrics (and pseudometrics) are closely connected to *norms*, which are a source of such natural measurements. Where metrics mathematise distances between points, a norm mathematises the *size* of a point (as a distance from an ‘origin’). For example, norms of the transition probability matrix of a discrete-time Markov chain measure how dynamic the chain is (Jafry and Schuermann, 2004). Similarly, pseudometrics such as Definition 3.5 yield a norm-like function on a set of matrices of real-valued functions.

#### Definition 6.4

Let *S* ⊆ ℝ^2^, *k ∈* ℕ, and *p, q* ∈ ℝ_*≥*1_. We define the norm-like function *n*_*p,q*_ : 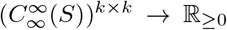 as follows: suppose 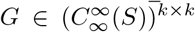.Let 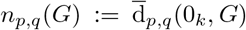 where 0_*k*_ is the *k*-by-*k* matrix of zeroes.

For ecosystem models, these norms are a way of measuring how dynamic an ecosystem is. Definition 6.4 yields a norm on *L*-ecosystems. Note that 0_*k*_ in Definition 6.4 is *not* an ecosystem (i.e. not in ℰ), as it violates either Axiom F. or G., but we can use it to define the norm. This yields a notion of the dynamical intensity of an ecosystem. An ecosystem with a large norm is very dynamic (as it is far from the 0_*k*_ matrix), whilst one with a smaller norm is less dynamic.

Some ecologically relevant measurements of the *L*-ecosystem are not continuous, and are therefore not natural measurements under our definition. For example, define # : *ℰ*→ℝ_*≥*0_ as #(*e*) is the number of non-zero entries in the matrix *G*^*e*^. Then #(*e*) is discontinuous. Nevertheless, counting the number of pairwise interactions in an ecosystem is the starting point for many measurements of food web structure (e.g. MacDonald, 1979).

### 6.4 Ecosytem Types and Connectedness

#### 6.4.1 Connected Spaces

There is a well-known historical controversy between those who believe in a natural classification of ecosystems (e.g. Clements, 1936, pp. 253, 282) and those who believe such classifications are “merely abstract extrapolations of the ecologist’s mind” (Gleason, 1926, p. 9). Simplified modern accounts (e.g. Begon et al., 2006, section 16.3.3) may overstate the extent of disagreement among participants (Hagen, 1992, chapter 2), but the controversy remains an important conceptual issue. We can address this question mathematically using connectedness. Our approach applies to any continuous ℝ-valued measurement on ecosystems, such as the norm-like function, *n*_*p,q*_ from Definition 6.4. A connected set has ‘one part’, as made precise in Theorem 6.5. In general, a proposed measurement of ecosystems, *m*, has two possible types of range, Ran(*m*):

GLEA. *m* classifies ecosystems into exactly one type, i.e. Ran(*m*) is an interval, so that if *m*(*e*) *< m*(*f*) are ecosystem *m*-values, and *m*(*e*) *< v < m*(*f*), then there is some ecosystem *g* such that *m*(*g*) = *v*, or

CLEM. *m* classifies ecosystems into more than one type, i.e. Ran(*m*) has disconnected parts (for example [0, 1] and [2, 3], so that no ecosystem takes an *m* value such as 1.5 in the space between these parts).

The main result is as follows:

##### Theorem 6.5.

Let *p, q ∈* ℝ_*≥*1_. Then *ℰ* is *τ*_*E,p,q*_-connected. So, if *m* : *ℰ →* ℝ is a *τ*_*E,p,q*_-continuous function, then *m*(*E*) is an interval *I ⊆* ℝ. A similar result holds if *p* = *∞* or *q* = *∞*.

Note that a function 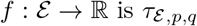-continuous if the preimage of an open set in ℝ is a union of balls, 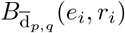, where *r*_*i*_ *>* 0.

So, if *m* is continuous, then GLEA. holds: *any* (continuous) measurement of *L*-ecosystems yields only one type. Under our axioms, it is possible to continuously deform any *L*-ecosystem to any other: for any choice of a small distance *δ >* 0, we can deform *e* into *f* by moving through intermediate *L*-ecosystems *e*_*i*_ such that *d*(*e*_*i*_, *e*_*i*+1_) *< δ* each time.

Hence, if *m* is a continuous measurement, then *m* exhibits the intermediate value theorem on ℰ (see Rudin, 1964, Theorems 4.22 and 4.23). This has far-reaching implications. For example, suppose we measure the norm of the matrix *G* of interspecific functions, or any other continuous real-valued function *m* of this matrix, on *L*-ecosystems. As *m* is continuous, if we have *L*-ecosystems *e, e*′ such that *m*(*e*) *< m*(*e*′), then for any *x* ∈ (*m*(*e*), *m*(*e*′)), there is an *L*-ecosystem *e*^*′′*^ such that *m*(*e*^*′′*^) = *x*.

Discontinuous measurements split the space of *L* ecosystems into different types when the measurement ‘builds in’ or ‘pre-categorises’ *L*-ecosystems into distinct types, e.g. the measurement #() from Section 6.3.

#### 6.4.2 Disconnected Spaces

The story is more complicated if the underlying set of ecosystems changes. Theorem 6.5 relies on the ecological axioms of Definition 6.1. A slight change to what counts as an ecosystem may yield a disconnected space of models, and hence distinct ecosystem types. Here we outline possible examples based on changing the ecological mechanisms between species. Firstly, we allow a mix of competition and mutualism. Secondly, we allow only mutualism or only competition. For readability, our description is high level: the details resemble *ℰ*, and we explain where they differ.

##### Example 6.6.

Suppose we require each producer, *i*, to have *g*_*i,i*_ *>* 0, and allow only competition (i.e. *g*_*i,j*_, *g*_*j,i*_ *<* 0) and mutualism (i.e. *g*_*i,j*_, *g*_*j,i*_ *>* 0). Call the resulting space of ecosystems *ℰ*′. Let 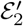 be the *e ∈ ℰ*′ with exactly two species. Note that any 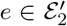 exhibits only mutualism or competition between the two distinct species. Then 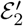 is not connected (e.g. one can put any mutualistic ecosystem in a small ball containing only mutualistic ecosystems). So, with a suitable notion of measurement, 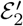 forms two separated types, corresponding to whether the species are mutualistic or competitive.

Similarly if each ecosystem in a collection exhibits only mutualism or only competition, but not both, then a corresponding disconnectedness result holds. A parallel result holds for the space of ecosystems where ecosystems exhibit either predation only or mutualism only (or competition only).

However, this example is unsatisfying as a mathematical defence of ecosystem types. Each time, the types are explicitly coded in: the mutualistic category and the competitive category. Moreover, the space is divided into exactly two parts: the ‘mutualistic only’ part is connected (as in Theorem 6.5), as is the ‘competitive only’ part.

##### Question 6.7.

Is the space of ecosystems which can exhibit a mixture of both mutualism and competition connected?

## 7 Discussion

Here, we showed that there is at least one natural way to compare the dynamical properties of ecosystem models with different sets of species. Our approach involves mapping each species in the smaller of a pair of ecosystems to a unique species in the larger ecosystem, and measuring distances between the functions describing pairwise interactions. Below, we discuss how our approach relates to conceptually similar methods for statistical analysis of unlabelled graphs (Calissano et al., 2024), and suggest some possible alternative approaches. We showed that our approach can be used in practice, through two proof-of-concept case studies. In a set of models for a marine sessile community with different biological mechanisms (Boughton et al., 2023), one of the models had clearly distinct dynamical properties from all the others. In the enaR database of ecosystem models (Borrett and Lau, 2014), freshwater and marine models were somewhat distinct, and different versions of the same ecosystem model were sometimes very different. We suggest potential avenues for exploration of these results. We showed that for a set of axioms designed to capture the essential properties of predator-prey and producer systems, there is only one distinct kind of ecosystem under natural measurements of dynamical properties. This result has implications for the nature and classification of ecosystems.

Our approach can be thought of as a generalization of recent methods for statistical analysis of unlabelled graphs (Calissano et al., 2024). In those methods, graphs typically represent interactions between objects, such as grooming interactions among baboons within a group. Vertices are treated as unlabelled because they represent different sets of objects of the same kind in different graphs. Edges are associated with scalar- or vector-valued attributes (in a Euclidean space) representing some aspect of interaction (such as the frequencies with which pairs of baboons interact). Given a metric on the space of labelled graphs that makes use of these attributes, a pseudometric can be computed on the space of unlabelled graphs by minimizing over permutations of vertices. The only key difference in our approach is that the “attributes” representing interactions between species are functions in an infinite-dimensional Hilbert space (Rynne and Youngson, 2008, Definition 3.23) rather than a Euclidean space. Thus it may be possible to extend our approach to statistical procedures such as computing ecosystem-valued means and principal components (c.f. Calissano et al., 2024). Conversely, there are many situations in which ecologists can obtain data on the signs of pairwise interactions (Puccia and Levins, 1985, pp. 5-6, 127-139) or the rates of flow of energy or matter at equilibrium (Baird et al., 2004b, p. 806), but not the functions determining these quantities. In such situations, the methods described in Calissano et al. (2024) are a natural choice. There may also be situations in which ways of matching species between ecosystems other than the injective mapping we used are more appropriate. For example, some ecosystem models may contain aggregated taxa such as “carnivores” in Figure 4, while others may attempt to resolve all taxa to species level. This is related to the problem of clone-consistency (Ansmann and Bollenbach, 2021). It may sometimes be possible to choose a meaningful level of aggregation, such as the trophic species level (Briand and Cohen, 1984). Alternatively, one could match every species in a larger ecosystem in which taxa are not aggregated with exactly one in a smaller ecosystem in which taxa are aggregated. The mapping between species would not then necessarily be either injective or surjective. The right approach may depend on the particular case.

We showed that our approach can be used in two proof-of-concept case studies, marine sessile community models (Boughton et al., 2023) and the enaR database (Borrett and Lau, 2014), but we would benefit from more information about dynamics and more systematic sampling of ecosystems. In the marine sessile study, we fitted a range of parametric models and found systematic differences in their dynamics, but none was an entirely adequate description of the experimental data. Although some aspects of the biology of interactions between jellyfish polyps and other sessile marine organisms are well understood (Fernández-Alías et al., 2024), this knowledge may not be sufficient to specify an adequate model from first principles. In the enaR database, the assumptions we made to get from flow networks to interaction functions are rooted in ecological theory, but are nevertheless strong, and the resulting functions are unlikely to capture behaviour far from the state at which flows were estimated. In particular, assuming Lotka-Volterra consumer-resource interactions may be misleading when integrating over a wide range of abundances. In both cases, the obvious direction for improvement is better estimation of the functions describing interspecific interactions. A variety of methods exist for non-parametric estimation of these functions (e.g. Jost and Ellner, 2000; Bonnaffé and Coulson, 2023), which could be further developed to incorporate axioms constraining their forms. However, these methods require large amounts of data, which may limit the potential for application to large numbers of ecosystems. Furthermore, existing data on ecological interactions are unlikely to be a representative sample of the world’s ecosystems (Poisot et al., 2021). It would therefore be unwise to draw strong conclusions about systematic differences between habitat types until better data are available, even though there is good reason to expect such differences (Steele et al., 2019).

With our chosen axioms for predator-prey and producer ecosystems, we showed that there is only one kind of ecosystem under any natural measurement of dynamical properties. This is consistent with the majority view that in physical space, “One association merges gradually into the next without any apparent transition zone” (e.g. Gleason, 1926, p. 14) and does not support the idea that there is a natural classification of ecosystems (e.g. Clements, 1936, pp. 253, 282). Nevertheless, we believe that the superficial summary of Clements’ views presented to generations of undergraduate ecologists (e.g. Begon et al., 2006, p. 478) has inhibited progress in understanding the nature of ecosystems as entities for which dynamical processes, rather than abundances or species identities, are the fundamental properties. There are also important qualifications to our result. First, we focused on systems containing predation and production, but also showed that the set of mutualistic and competitive ecosystems can be disconnected. However, a single real ecosystem may have predation, production, competition and mutualism, and a full treatment will therefore require more general axioms. Our current axioms are also bio-sociological (focusing on interactions among organisms) rather than biogeochemical (focusing on transfer of matter or energy) in the sense of Hutchinson and Wollack (1940), and should in future include functions representing dynamics of non-living components. Second, even if the set *ℰ* of ecosystems is connected, not all members of this set may be possible in the real world. For example, if only some parts of an environmental gradient *l* between Earth and another planet can support life, then the ecosystems that actually occur on these planets could in practice be distinct (Figure 6) even if they are governed by common ecological laws (Jones, 2001; Damuth and Ginzburg, 2025, pp. 49, 125). Less extreme versions of this argument could operate even on a single planet. Third, selection at the ecosystem level (Borrelli et al., 2015) f for properties such as stability (Dunbar, 1960) or total energy flux (Lotka, 1922) might eliminate some ecosystems, leaving a disconnected set. Earlier arguments for selection of this kind are not widely accepted (Månsson and McGlade, 1993), but there is the potential for a non-adaptive form of selection to operate on communities or ecosystems without requiring distinct kinds of ecosystem (Damuth and Ginzburg, 2025, pp. 54-58), and to be an important source of ecological laws (Damuth and Ginzburg, 2025, pp. 47-51). We are not asserting the sorites paradox that because all ecosystems are of one kind, they are all the same under any natural measurement of dynamical properties. Clearly, the value of a natural measurement (which we define as being continuous) can differ between ecosystems, but a natural classification cannot be based on a continuous function whose domain is connected. The sorites paradox in its classical form (Barnes, 1982, pp. 33-34) is that because one grain of wheat does not form a heap, and adding a single grain to a collection of grains that is not a heap does not turn it into a heap, there is no such thing as a heap of wheat grains. Avoiding such a paradox in our case requires at least one of three things (Dzhafarov and Dzhafarov, 2010). First, an ecosystem could be of more than one kind (because the kinds are not well defined, or for other reasons). For example, there is a widespread belief that savannas and forests are distinct kinds of ecosystem (Ratnam et al., 2011), yet relying on vague terms such as “dominant”, “relatively shorter” and “no or few” to classify them (Ratnam et al., 2011, Key 1, Table 1) makes the distinction ill-defined. Second, for at least one ecosystem, there may be other ecosystems sufficiently close to it with different values of the natural measurement. This is likely to be the case for most non-constant natural measurements. Third, we may have a measurement that breaks up the set of ecosystems into equivalence classes with the same value of the measurement, for example the subsets of ecosystems for which the equilibrium with all species absent is locally stable or not, and the subsets of coral reefs with and without the potential for positive calcium carbonate production (Andersson, 2015). Such measurements are potentially of interest, but they are not natural measurements under our definition because they are not continuous, and they predefine a classification of ecosystems.

**Figure 6:**
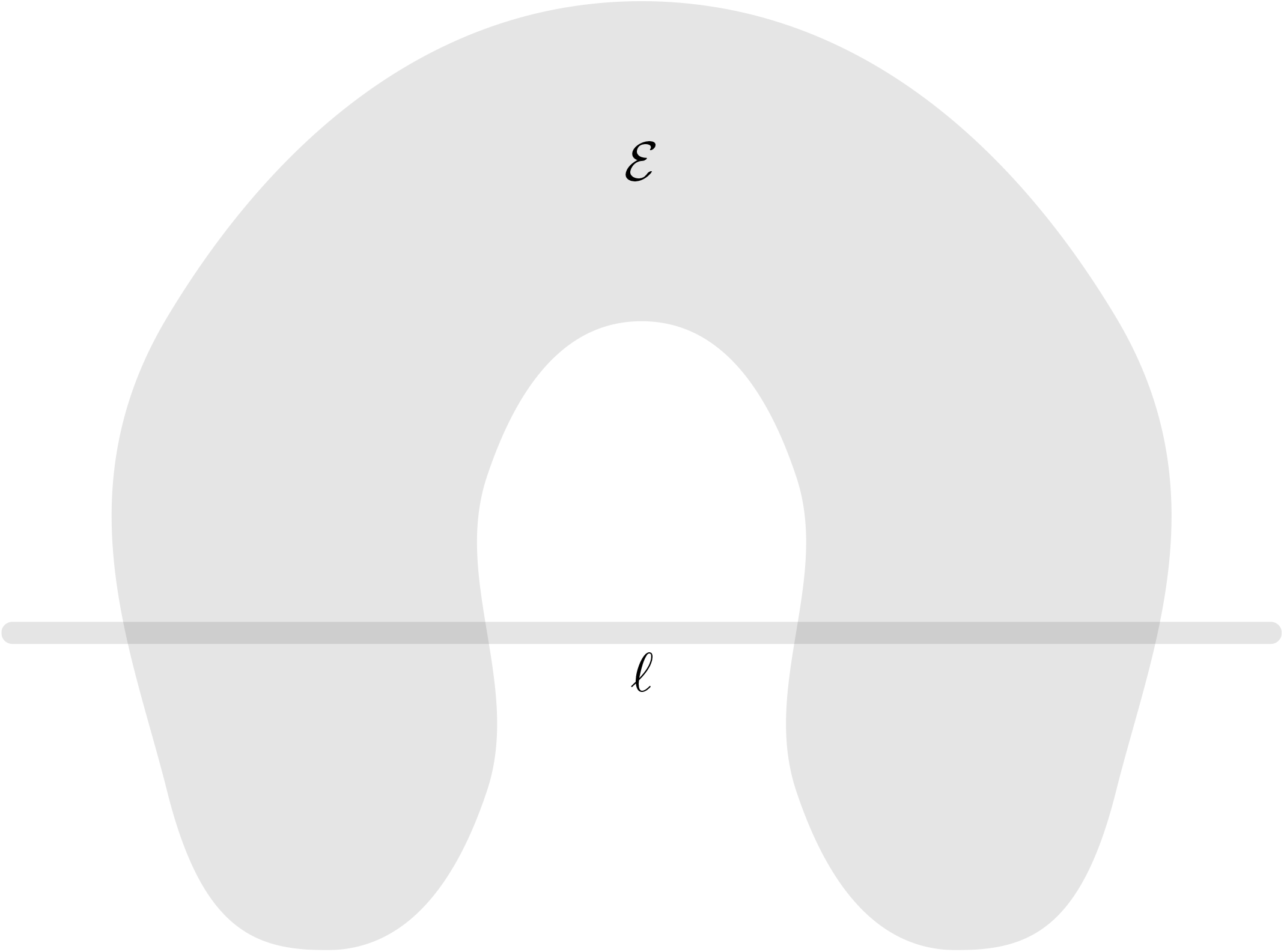
Even if the set *ℰ*of all possible ecosystems is connected, the subset *ℰ*∩*l* that occurs in nature along a gradient *l* may be disconnected.

In conclusion, we have shown that focusing on differences in dynamics, rather than in species identities or abundances, leads to new answers to old questions about the natural classification of ecosystems. We have also shown that our approach can in principle be applied to numerical comparisons of particular sets of ecosystem models. We developed a set of ecologically-motivated axioms intended to describe the properties of predator-prey systems, under which we showed that there is only one kind of ecosystem under any natural measurement of dynamics. There is scope for further development of general axioms covering other biological mechanisms such as competition and mutualism.

## A Example 1: Other Pseudodistances

**Figure A1:**
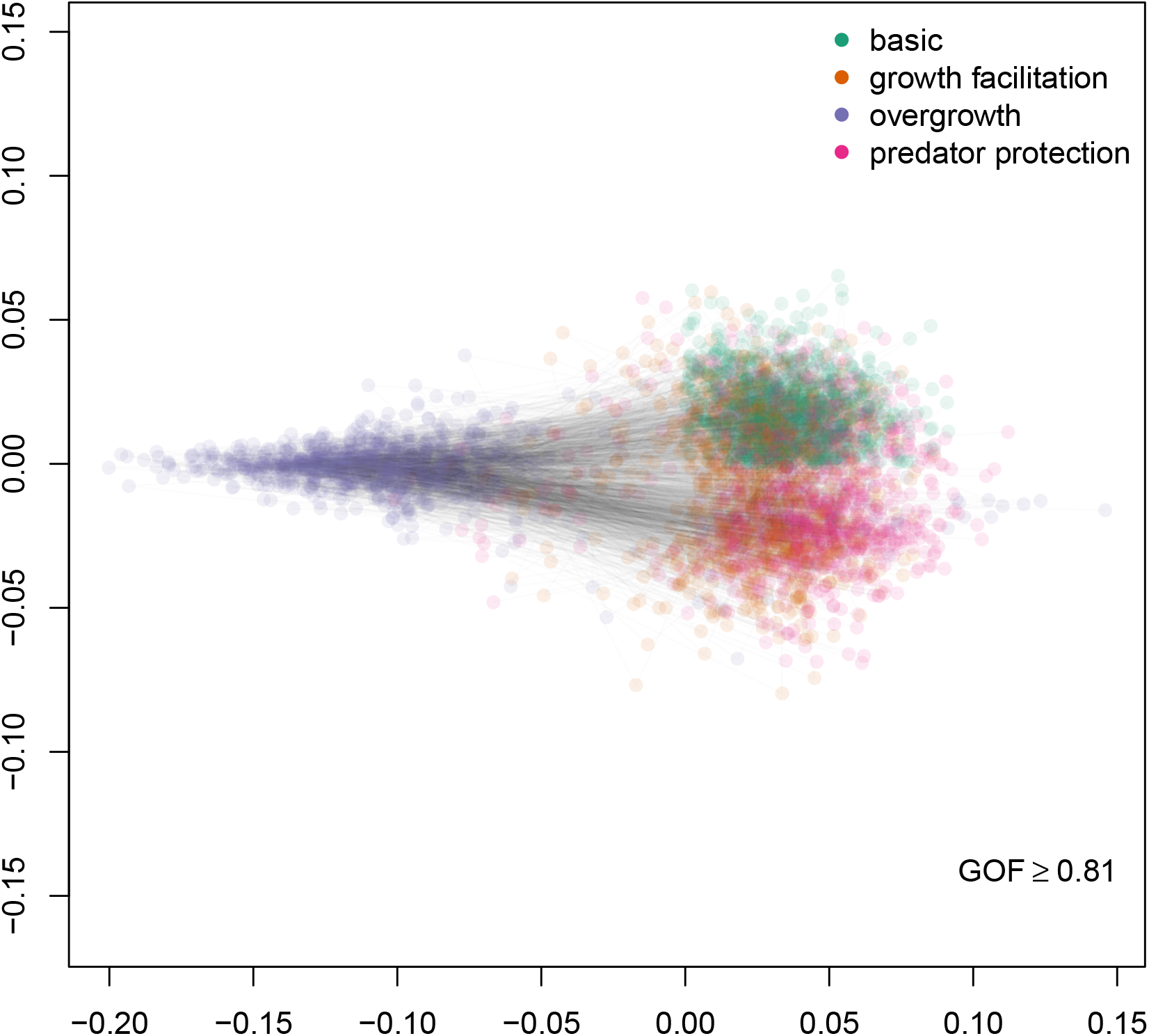
Classical multidimensional scaling of pseudodistances 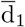 between 4 models for a sessile community. Each set of 4 coloured points (green: basic, orange: growth facilitation, purple: overgrowth, pink: predator protection) connected by lines represents a single pseudodistance matrix with parameters drawn from their posterior distributions (1000 draws in total). Goodness-of-fit (GOF) for the two-dimensional representation was at least 0.81 in all draws.

**Figure A2:**
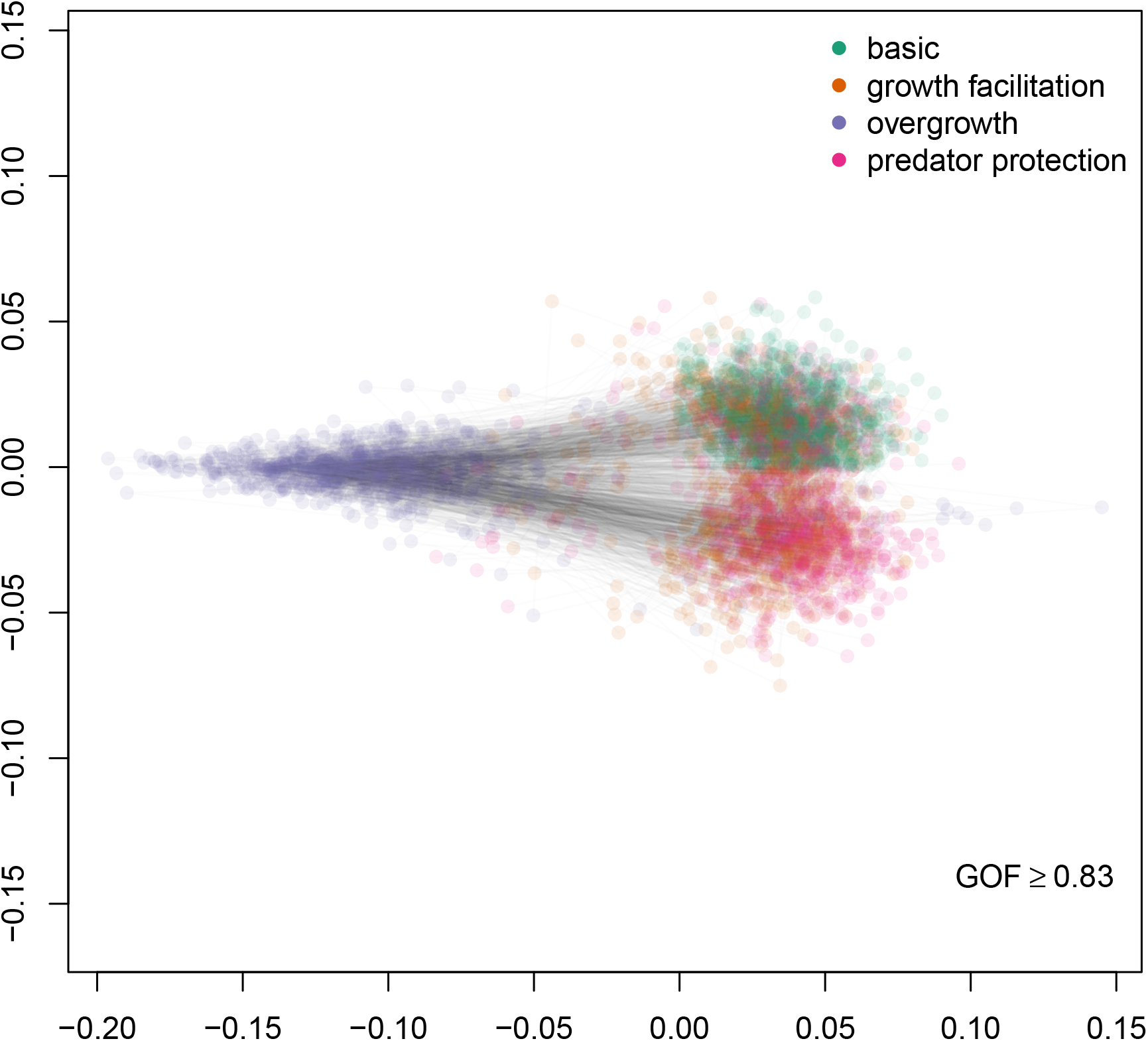
Classical multidimensional scaling of pseudodistances 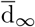 between 4 models for a sessile community. Each set of 4 coloured points (green: basic, orange: growth facilitation, purple: overgrowth, pink: predator protection) connected by lines represents a single pseudodistance matrix with parameters drawn from their posterior distributions (1000 draws in total). Goodness-of-fit (GOF) for the two-dimensional representation was at least 0.85 in all draws.

**Table A1:** Pseudodistances 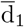 between 4 models for a sessile community (basic, growth facilitation, overgrowth, predator protection). Posterior means from 1000 draws, with 0.025− and 0.975− quantiles in brackets.

|  | basic | growth facilitation | overgrowth | predator protection |
| --- | --- | --- | --- | --- |
| basic | 0.00 (0.00, 0.00) | 0.05 (0.02, 0.09) | 0.14 (0.07, 0.23) | 0.05 (0.02, 0.10) |
| growth facilitation | 0.05 (0.02, 0.09) | 0.00 (0.00, 0.00) | 0.14 (0.07, 0.22) | 0.06 (0.02, 0.11) |
| overgrowth | 0.14 (0.07, 0.23) | 0.14 (0.07, 0.22) | 0.00 (0.00, 0.00) | 0.15 (0.07, 0.24) |
| predator protection | 0.05 (0.02, 0.10) | 0.06 (0.02, 0.11) | 0.15 (0.07, 0.24) | 0.00 (0.00, 0.00) |

**Table A2:** Pseudodistances 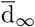 between 4 models for a sessile community (basic, growth facilitation, overgrowth, predator protection). Posterior means from 1000 draws, with 0.025− and 0.975 − quantiles in brackets.

|  | basic | growth facilitation | overgrowth | predator protection |
| --- | --- | --- | --- | --- |
| basic | 0.00 (0.00, 0.00) | 0.10 (0.03, 0.22) | 0.37 (0.17, 0.58) | 0.11 (0.04, 0.22) |
| growth facilitation | 0.10 (0.03, 0.22) | 0.00 (0.00, 0.00) | 0.38 (0.18, 0.58) | 0.12 (0.04, 0.24) |
| overgrowth | 0.37 (0.17, 0.58) | 0.38 (0.18, 0.58) | 0.00 (0.00, 0.00) | 0.37 (0.18, 0.57) |
| predator protection | 0.11 (0.04, 0.22) | 0.12 (0.04, 0.24) | 0.37 (0.18, 0.57) | 0.00 (0.00, 0.00) |

## B Example 2: other pseudodistances, scree plot and data sources

**Figure A3:**
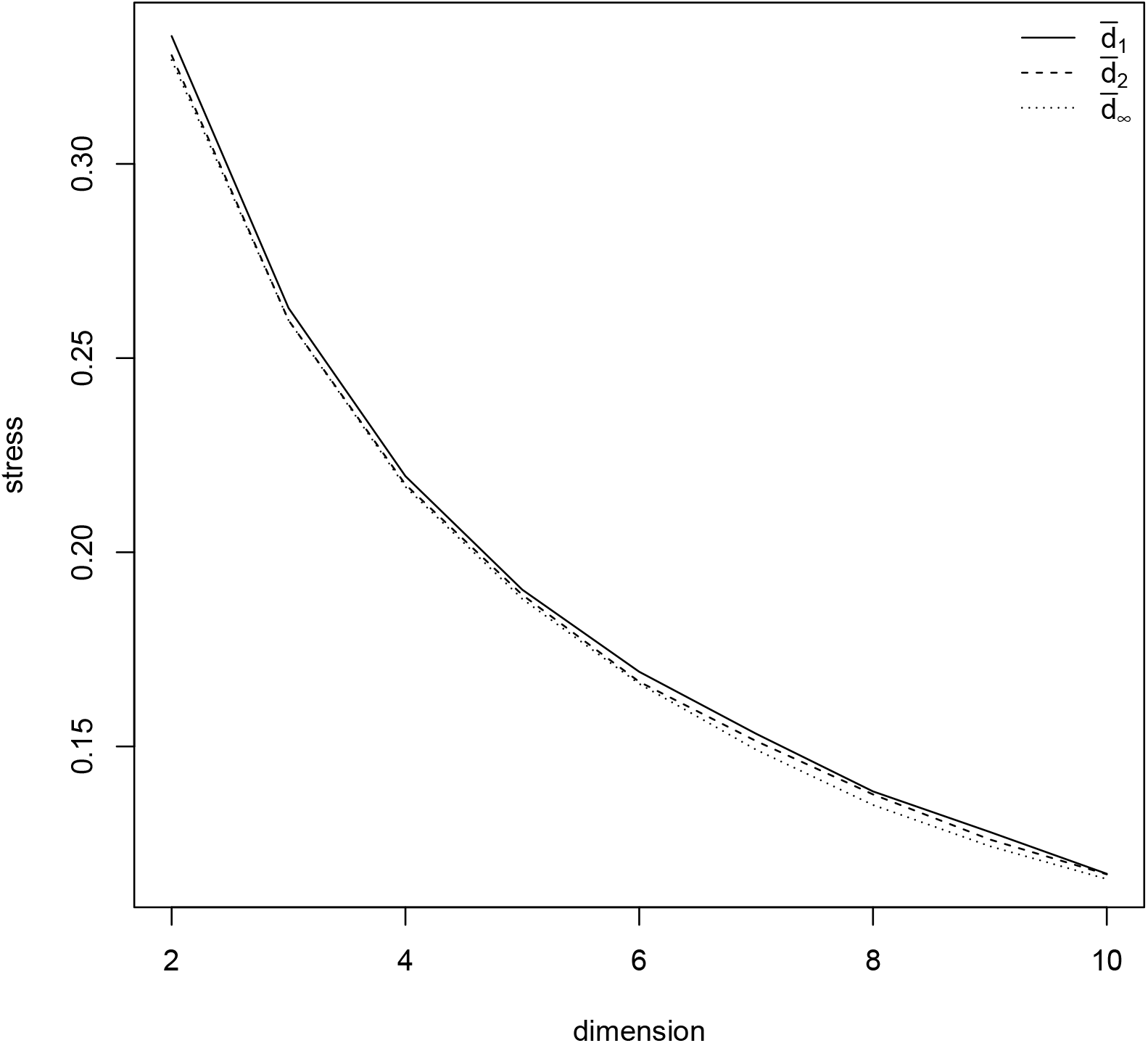
Relationship between stress and dimension for non-metric multidimensional scaling of pseu-dodistances 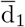 (solid line), 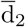 (dashed line) and 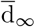 (dotted) among 50 ecosystem models from the troModels compilation in the enaR package (Borrett and Lau, 2014).

**Figure A4:**
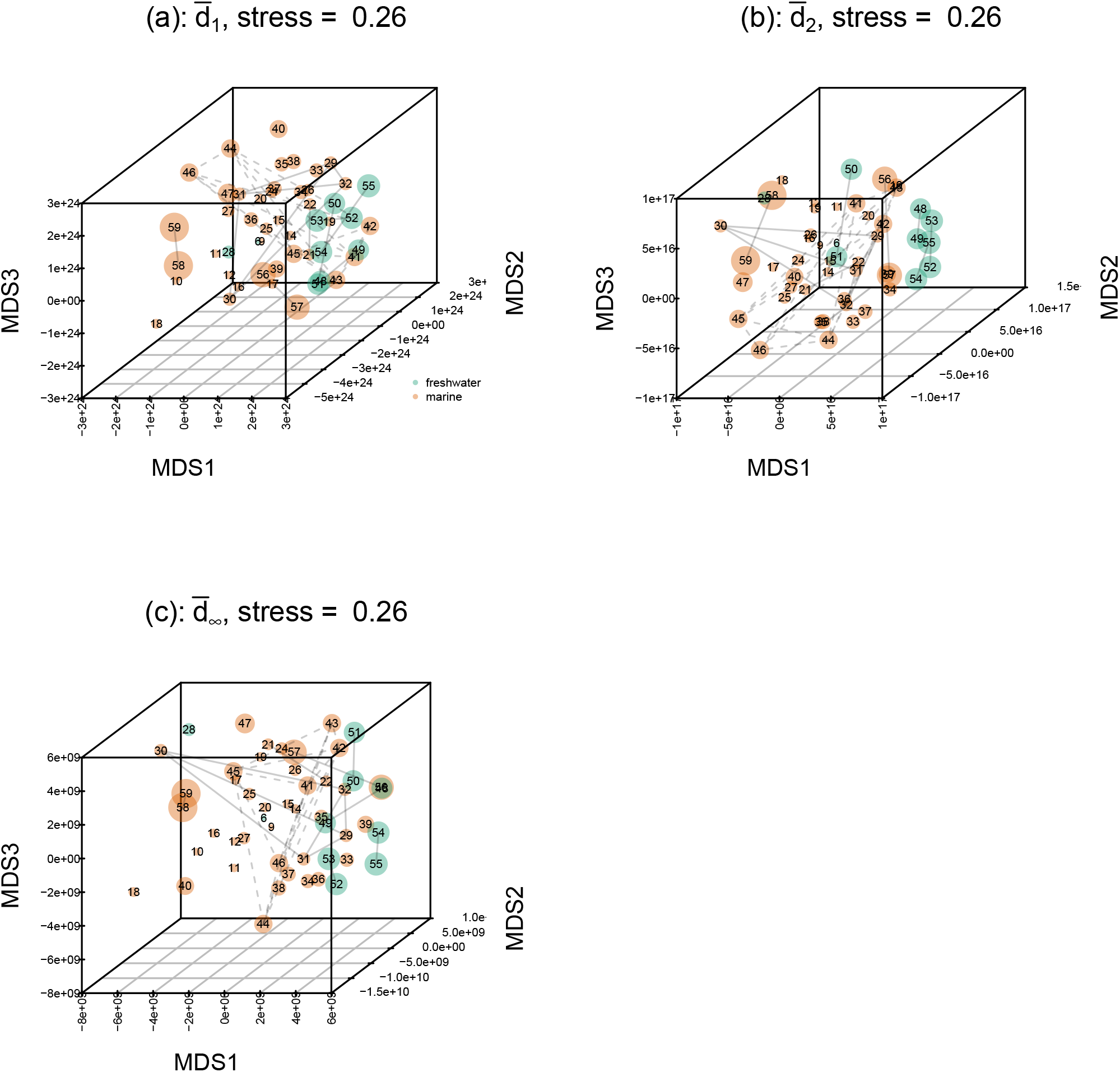
Three-dimensional non-metric multidimensional scaling of pseudodistances 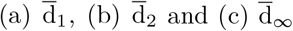 among 50 ecosystem models from the troModels compilation in the enaR package (Borrett and Lau, 2014). Numbers on the ordinations correspond to model numbers in troModels. The stress is indicated for each ordination. Point volumes are proportional to the number of living compartments in the model. Point colours represent habitat types (green freshwater, orange marine). Lines connect models for different versions of the same ecosystem such as sites or times. Dashed lines indicate the large cluster of St Marks Seagrass ecosystems at different sites and times, numbers 41 to 46 (Baird et al., 1998).

**Table A3:**
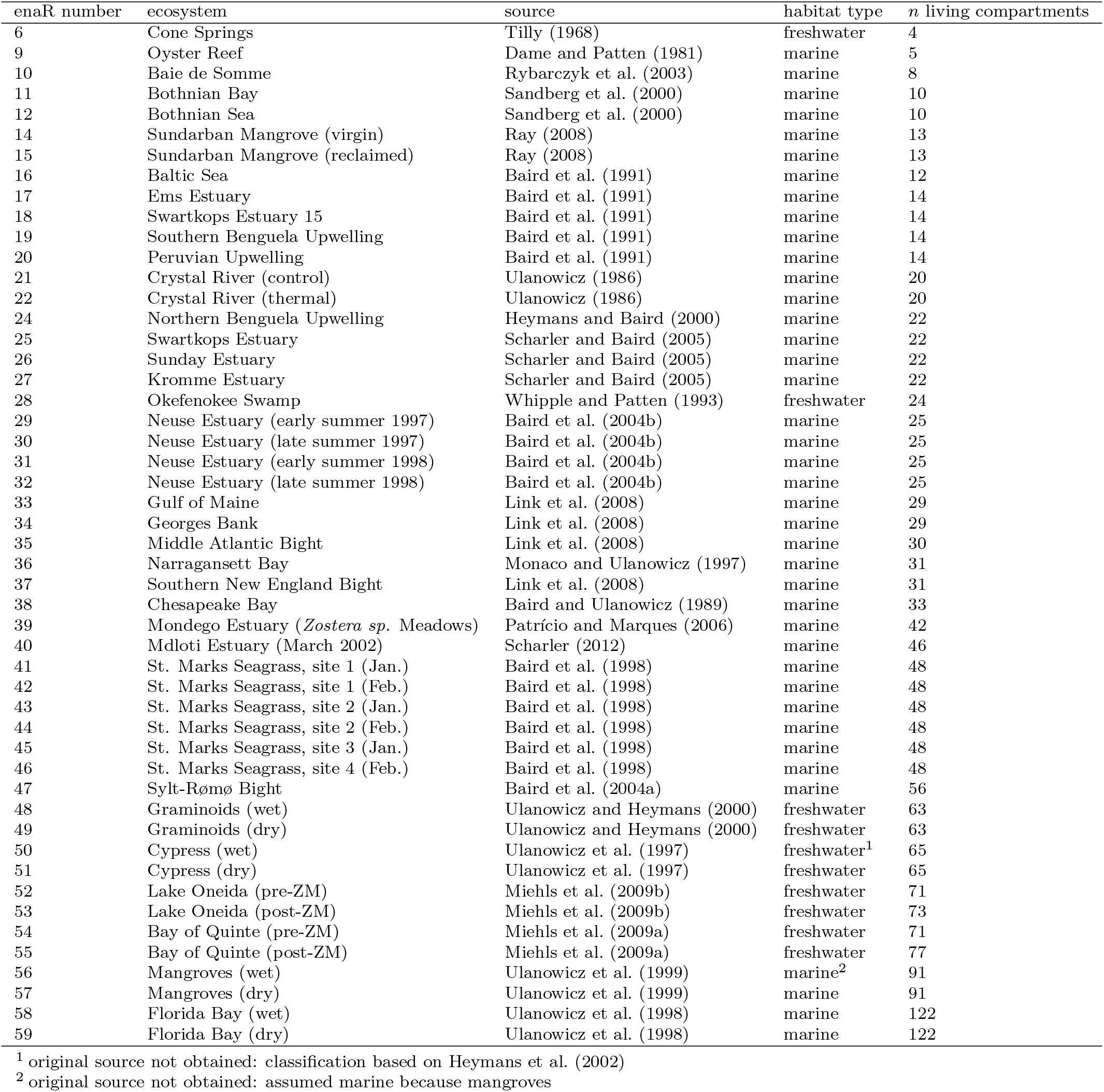
Data sources for the enaR troModels example. Columns: enaR number and ecosystem refer to model numbers, names and original sources (Borrett and Lau, 2014); habitat type classified as described in the main text, section 5.3 (based on original sources except as indicated below); *n* living compartments obtained by counting the number of `TRUE` values in the `living` variable for the model in the `troModels` list in `enaR`.

| enaR number | ecosystem | source | habitat type | $n$ living compartments |
| --- | --- | --- | --- | --- |
| 6 | Cone Springs | Tilly (1968) | freshwater | 4 |
| 9 | Oyster Reef | Dame and Patten (1981) | marine | 5 |
| 10 | Baie de Somme | Rybarczyk et al. (2003) | marine | 8 |
| 11 | Bothnian Bay | Sandberg et al. (2000) | marine | 10 |
| 12 | Bothnian Sea | Sandberg et al. (2000) | marine | 10 |
| 14 | Sundarban Mangrove (virgin) | Ray (2008) | marine | 13 |
| 15 | Sundarban Mangrove (reclaimed) | Ray (2008) | marine | 13 |
| 16 | Baltic Sea | Baird et al. (1991) | marine | 12 |
| 17 | Ems Estuary | Baird et al. (1991) | marine | 14 |
| 18 | Swartkops Estuary 15 | Baird et al. (1991) | marine | 14 |
| 19 | Southern Benguela Upwelling | Baird et al. (1991) | marine | 14 |
| 20 | Peruvian Upwelling | Baird et al. (1991) | marine | 14 |
| 21 | Crystal River (control) | Ulanowicz (1986) | marine | 20 |
| 22 | Crystal River (thermal) | Ulanowicz (1986) | marine | 20 |
| 24 | Northern Benguela Upwelling | Heymans and Baird (2000) | marine | 22 |
| 25 | Swartkops Estuary | Scharler and Baird (2005) | marine | 22 |
| 26 | Sunday Estuary | Scharler and Baird (2005) | marine | 22 |
| 27 | Kromme Estuary | Scharler and Baird (2005) | marine | 22 |
| 28 | Okefenokee Swamp | Whipple and Patten (1993) | freshwater | 24 |
| 29 | Neuse Estuary (early summer 1997) | Baird et al. (2004b) | marine | 25 |
| 30 | Neuse Estuary (late summer 1997) | Baird et al. (2004b) | marine | 25 |
| 31 | Neuse Estuary (early summer 1998) | Baird et al. (2004b) | marine | 25 |
| 32 | Neuse Estuary (late summer 1998) | Baird et al. (2004b) | marine | 25 |
| 33 | Gulf of Maine | Link et al. (2008) | marine | 29 |
| 34 | Georges Bank | Link et al. (2008) | marine | 29 |
| 35 | Middle Atlantic Bight | Link et al. (2008) | marine | 30 |
| 36 | Narragansett Bay | Monaco and Ulanowicz (1997) | marine | 31 |
| 37 | Southern New England Bight | Link et al. (2008) | marine | 31 |
| 38 | Chesapeake Bay | Baird and Ulanowicz (1989) | marine | 33 |
| 39 | Mondego Estuary (*Zostera sp.* Meadows) | Patrício and Marques (2006) | marine | 42 |
| 40 | Mdloti Estuary (March 2002) | Scharler (2012) | marine | 46 |
| 41 | St. Marks Seagrass, site 1 (Jan.) | Baird et al. (1998) | marine | 48 |
| 42 | St. Marks Seagrass, site 1 (Feb.) | Baird et al. (1998) | marine | 48 |
| 43 | St. Marks Seagrass, site 2 (Jan.) | Baird et al. (1998) | marine | 48 |
| 44 | St. Marks Seagrass, site 2 (Feb.) | Baird et al. (1998) | marine | 48 |
| 45 | St. Marks Seagrass, site 3 (Jan.) | Baird et al. (1998) | marine | 48 |
| 46 | St. Marks Seagrass, site 4 (Feb.) | Baird et al. (1998) | marine | 48 |
| 47 | Sylt-Rømø Bight | Baird et al. (2004a) | marine | 56 |
| 48 | Graminoids (wet) | Ulanowicz and Heymans (2000) | freshwater | 63 |
| 49 | Graminoids (dry) | Ulanowicz and Heymans (2000) | freshwater | 63 |
| 50 | Cypress (wet) | Ulanowicz et al. (1997) | freshwater<sup>1</sup> | 65 |
| 51 | Cypress (dry) | Ulanowicz et al. (1997) | freshwater | 65 |
| 52 | Lake Oneida (pre-ZM) | Miehls et al. (2009b) | freshwater | 71 |
| 53 | Lake Oneida (post-ZM) | Miehls et al. (2009b) | freshwater | 73 |
| 54 | Bay of Quinte (pre-ZM) | Miehls et al. (2009a) | freshwater | 71 |
| 55 | Bay of Quinte (post-ZM) | Miehls et al. (2009a) | freshwater | 77 |
| 56 | Mangroves (wet) | Ulanowicz et al. (1999) | marine<sup>2</sup> | 91 |
| 57 | Mangroves (dry) | Ulanowicz et al. (1999) | marine | 91 |
| 58 | Florida Bay (wet) | Ulanowicz et al. (1998) | marine | 122 |
| 59 | Florida Bay (dry) | Ulanowicz et al. (1998) | marine | 122 |

<sup>1</sup> original source not obtained: classification based on Heymans et al. (2002)
<sup>2</sup> original source not obtained: assumed marine because mangroves

## C Generative Axioms

In Definition 6.1, we defined the class of *L*-ecosystems. We first identified a class of candidates models, then specified which were *L*-ecosystems. The other axiomatic approach is to ‘build’ ecosystems out of certain ‘basic’ ecosystems (cf. building sets in set theory using `ZFC,` see (Jech, 2003, Chapter 1)). Here, we state suitable *generative* axioms from which we can build ecosystems. We define the class of *G*-ecosystem as the ecosystem models which are thus built. Notably, this class coincides with the class of *L*-ecosystems.

We first define a *dynamics assignment function* which describes the pairwise dynamics for each pair of species in an ecosystem: different ecosystem models can have the same dynamics assignment function (e.g. if one ecosystem includes another), and an ecosystem model is compatible with multiple dynamics assignment functions. The dynamics assignment function takes a pair of natural numbers, representing (the labels of) two species, to three pieces of data:

I. the conversion of *i* consumed by *j* into units of *i* in the case of predation (*c*_*i,j*_ below),
II. the predation relation (*P*_*i,j*_ below), and
III. the contribution to the proportional growth rate of *n* with respect to *m* (*g*_*i,j*_ below).^1^

### Definition C.1.

Let 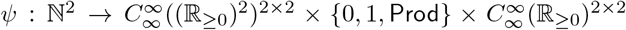. We define 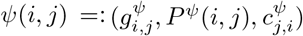. When *ψ* is clear from context, we denote these by *g*_*i,j*_, *P* (*i, j*), and *c*_*i,j*_. We say *ψ* is a *dynamics assignment function* if and only if all of the following hold for any *i, j ∈* ℕ, and for all *x, y ∈* ℝ_*≥*0_.

1. (Prey Consumption and Predator Growth) If *P* (*i, j*) = 1, then *g*_*i,j*_(*x, y*) *>* 0 *> g*_*j,i*_(*x, y*).
2. (Only Predation and Production) If *P* (*i, j*) = 0 and *i ≠ j*, then *g*_*i,j*_(*x, y*) = 0.
3. (Non-producers need Prey) If *P* (*i, i*) *≠* Prod, then *g*_*i,i*_(*x, x*) *<* 0.
4. (Producers are Self-Limiting) If *P* (*i, i*) = Prod then *g*_*i,i*_(*x, x*) is strictly decreasing in *x*.
5. (Monotonicity) if *P* (*i, j*) = 1 then *g*_*i,j*_(*x, y*) is strictly increasing in *y* and *g*_*j,i*_(*x, y*) is strictly decreasing in *x*.
6. (Inefficient Conversion) If *P* (*i, j*) = 1 then 0 *≤ c*_*i,j*_(*x*) *< x*, otherwise *c*_*i,j*_ = 0.

Next, we define *G*-ecosystems. A *G*-ecosystem consists of a finite sequence of natural numbers (with no repeats), representing the labels of some species, plus some information extracted from a fixed dynamics assignment function, *ψ*. We say *e* is *founded* on *ψ*.

### Definition C.2.

Let *S* = (*s*_*i*_)_*i∈*|*S*|_ be a finite sequence of natural numbers, without repeats. Let *ψ* be a dynamics assignment function. Let *e* = (*S*^*e*^, *G*^*e*^, *P* ^*e*^, *c*^*e*^) where *P* ^*e*^ = {0, 1, `Prod`}^|*S*|*×*|*S*|^, and *G*^*e*^ and *c*^*e*^ are |*S*^*e*^|-by-|*S*^*e*^| matrices such that 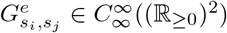 and 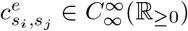 where *P* ^*e*^(*i, j*) = *P* ^*ψ*^(*s*_*i*_, *s*_*j*_), 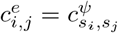, and 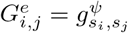.

Suppose in addition that *i ∈* |*S*|. Let 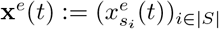 be such that 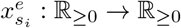 are solutions to the following |*S*|-many equations:

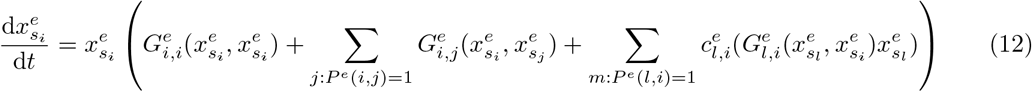

We call 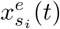 the *abundance* of species *s*_*i*_ at *t*. We say that *e* is an *G-ecosystem*, and write *e ∈ G*, if it satisfies the following condition for all *i, j* ∈ |*S*| and all *t ∈* ℝ_*≥*0_:

7. (Producers can Increase when Everything is Rare) If 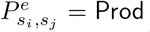 then *i* = *j* and for all *k ∈* |*S*|, there exist abundances *b*_*k*_ *∈* ℝ_*≥*0_, such that if 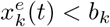, then 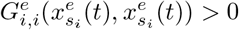.

We say that *e* is *founded* by *ψ*, and that *S* is the *sequence of species (labels)* of *e*. As a shorthand, we can drop the *e* superscripts, and write 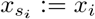 (though really *x*_*i*_ represents the *i*^th^ species, not the 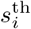 species).

We now state the result showing how to build all *G*-ecosystems from the basic ecosystem with only two species. The building methods are (Permuting), where the species-label are permuted, (Shrinking), where we remove a species from an ecosystem, and (Gluing), where we combine ecosystems.^2^

If (*s*_*n*_)_*n∈N*_, (*t*_*m*_)_*m∈M*_ are sequences of natural numbers with no repeats, let merge((*s*_*n*_)_*n∈N*_, (*t*_*m*_)_*m∈M*_) be the sequence (with no repeats) consisting of (*s*_*n*_)_*n∈N*_ followed by all entries of the set *{t*_*m*_ : *m ≤ M}\{s*_*n*_ : *n ≤ N}* ordered by the ordering of (*t*_*m*_)_*m∈M*_.

### Theorem C.3.

The *G*-ecosystems are exactly the models resulting from the following constructions.

1. (Basic Ecosystems) Let *n, m ∈* ℕ. If there is a dynamics assignment function, *ψ*, such that 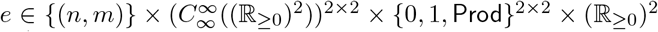 where *π*_2_(*e*) = *g*_*ψ*_(*n, m*) and *π*_3_(*e*) = *P* ^*ψ*^(*n, m*), then *e* is a *G*-ecosystem founded by *ψ*.
2. (Permuting) If *e* ∈ *G* (founded by *ψ*), and *π*_1_(*e*) = *s*, a sequence of natural numbers, and *σ* : *s* → *s*′ is a permutation of *s*, then the model resulting from *σ*-permuting the indices in all entries in *e*, which we denote *σ*(*e*), is a *G*-ecosystem (founded by *ψ*).
3. (Shrinking) Let *e* = (*S, G, P, c*) ∈ *G* (founded by *ψ*). Let *e*′ be the result of uniformly removing column *i* and row *i* from *e* from each of *S, G, P*, and *C*. If for all *t, x*_*i*_(*t*) = 0 then *e*′ ∈ *G* (and is founded by *ψ*).
4. (Gluing) If *e, f ∈ G* are both founded by *ψ*, then there is an *G*-ecosystem, denoted *e* + *f*, founded by *ψ*, such that *S*^*e*+*f*^ = merge(*S*^*e*^, *S*^*f*^), such that for all *i, j ∈ e, P* ^*e*+*f*^ (*i, j*) = *P* ^*e*^(*i, j*), *c*^*e*+*f*^ (*i, j*) = *c*^*e*^(*i, j*) and *g*^*e*+*f*^ = *g*^*e*^, and so too for *f*.

*Proof*. We first prove that if *e* is a model which is constructed using the five constructions, then *e* ∈ *G*. (Basic Ecosystems), (Permuting), and (Shrinking) are immediate from the definition of *G*. So we prove (Gluing).

For (Gluing), let *S*^*e*+*f*^ = merge(*S*^*e*^, *S*^*f*^). Let *e* + *f* be defined in the following way: for all *i, j ∈ S*^*e*+*f*^, 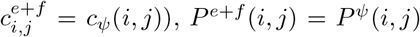 and 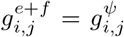. Then *e* + *f ∈ G*, is founded by *ψ*, and agrees with *e* and *f* on their common species, as required.

Conversely, we prove that if *e ∈ G*, then *e* can be constructed in this way. For this, let *e ∈ G* be founded by *ψ*. Let *π*_1_(*e*) = *S*, the sequence of species. For each pair *s*_*i*_, *s*_*j*_ *∈* Ran(*S*) (i.e. the *s*_*i*_ with *i ∈* |*S*|), let *e*_*i,j*_ be the ecosystem of size 2 founded by *ψ* which contains exactly the species *s*_*i*_ and *s*_*j*_ (possibly allowing *s*_*i*_ = *s*_*j*_). As Ran(*S*) is finite, we can partition Ran(*S*) into finitely many pairs {*s*_1_, *s*_2_}, …, {*s*_2*n*+1_, *s*_2*n*+2_}, where *n* depends on the size of *S*, with possibly one improper pair {*s*_|*S*|_, *s*_|*S*|_} if |*S*| is odd. By (Basic Ecosystems), each of these pairs constitutes a *G*-ecosystem. By applying (Gluing) finitely many times, for each of the corresponding *e*_*i,j*_ for each of these pairs, we recover *e*, i.e.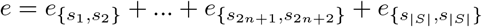.

There are two main differences between an *L*-ecosystem, *f*, and a *G*-ecosystem, *e*. Firstly, the vector of abundance functions, **x**(*t*), is explicit in *f* (as *π*_1_(*f*)), but implicit in *e* (as the solution of Equation 12). Secondly, the species of *f* are labelled by a *k* ∈ ℕ, whilst those of *e* are labelled by a non-repeating sequence of natural numbers (whose range need not be an initial segment of the order of ℕ, of the form 1, 2, 3, 4,..). For ecosystem models, the two definitions coincide:

### Theorem C.4.

Let *e ∈ G*. Then there is an *f ∈ ℰ* over a *k ∈* ℕ such that *e* and *f* have the same number of species, and for all *s*_*i*_, *s*_*j*_ *∈* Ran(*S*^*e*^), *P* ^*e*^(*s*_*i*_, *s*_*j*_) = *P* ^*f*^ (*i, j*), 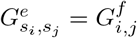, and 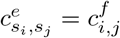.

Conversely, if *f ∈ ℰ* over *k ∈* ℕ, then there is an *e ∈ G*, with Ran(*S*^*e*^) = *k* (hence |*S*^*e*^| is finite), such that *e* and *f* have the same number of species, and for all *s*_*i*_, *s*_*j*_ ∈ `Ran` (*S*^*e*^), *P* ^*e*^(*s*_*i*_, *s*_*j*_) = *P* ^*f*^ (*i, j*), 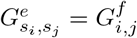, and 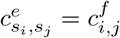.

*Proof*. We first show that any *G*-ecosystem, *e*, has a corresponding *L*-ecosystem. Let *e* be founded by *ψ*. Let *N* (= | `Ran` (*S*^*e*^)|) be the number of species in *e*, clearly *N* is an initial segment of ℕ, of the form 1, 2, 3, …. Choose any vector of starting values, **x**(0), such that 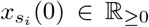. As *e ∈ G*, for every *i, j ∈* |*S*|, we have that 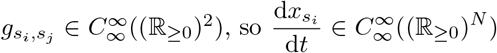 (from Equation 12). Hence, these form a set of differential equations with a solution **x**(*t*). It suffices to check that *T* (*e*) := (*G*^*e*^, **x**, *P* ^*e*^, *c*^*e*^) is an *L*-ecosystem. We check each of Axioms A. to I. in turn. The properties mostly follow from the corresponding properties of the dynamics assignment function, *ψ* or of *e* itself. Only Axioms A. and I., are not immediate. For these:

A. Equation 12 is a Kolmogorov equation 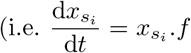 for some function *f*), so if 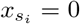, then 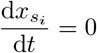. Hence, by the constant value theorem, 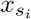 is constant, with value 0 for all future times. Hence *A*. holds.
I. This holds by the definition of *T* (*e*): we defined *T* (*e*) so that **x** is a solution to Equation 12, hence 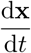 satisfies Equation 11 by basic differentiation.

Next, we prove the converse, that an *L*-ecosystem, *f*, has a corresponding *G*-ecosystem. Let **k** be the *k*-length sequence such that *s*_*i*_ = *i* (i.e. **k** = (1, …, *k*)). Let *R*(*f*) = (**k**, *G*^*f*^, *P* ^*f*^, *c*^*f*^). It suffices to check that *R*(*f*) is founded by some dynamic assignment function. We define a dynamic assignment function, *Q*(*f*), like so:

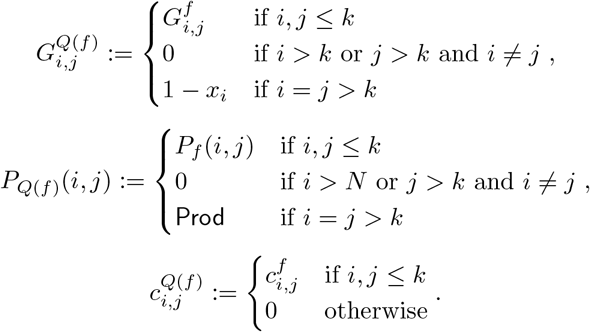

The third case of each of 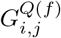 and *P*_*Q*(*f*)_ (*i, j*) are included so that the dynamics assignment function is well defined on the species not included in *e*. This amounts to adding a number of independent producers.

Clearly *Q*(*f*) founds *R*(*f*), so we check that *Q*(*f*) is a dynamic assignment function. The necessary properties of *ψ* can be checked easily, e.g. (Inefficient Conversion) for *Q*(*f*) follows from (Inefficient Conversion) for *f*. Finally, note that:

2. for *e* follows from (Producers can Increase when Everything is Rare) for *f* for *i ≤ k*, and if *i > N* then 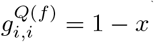 which satisfies (Producers can Increase when Everything is Rare).

## D Alternative Pseudometrics

Definition 3.2 builds in some uniformity: for each pair *i, j ≤ k*, the coordinate metric was constant, e.g. always the *L*_*p*_ metric. Moreover, each coordinate is equally weighted. More general definitions are possible, e.g. accounting for weighting:

### Definition D.1.

Let *q* = *∞* or *q ∈* ℝ_*≥*1_. For each pair *i, j ≤ k*, let *n*_*i,j*_ *∈* ℕ, 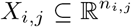, *I*_*i,j*_ *⊆* ℝ be an interval, *p*_*i,j*_ be such that *p*_*i,j*_ = *∞* or *p ∈* ℝ_*≥*1_, and let *c*_*d,w*_ *∈* ℝ_*≥*0_. We call *c*_*i,j*_ the *weighting* of the (*i, j*)^*th*^-coordinate. Let *G, G*′ be (*k × k*)-matrices such that 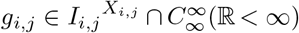, and likewise for 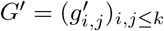. We define the following pseudometric:

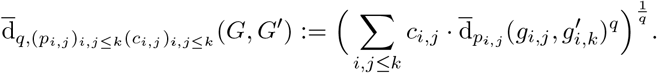

## E Proof Outlines

*Proof of Theorem 3.4*. We first consider 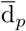. We check each of the three properties of a pseudometric. Using the identity bijection, note that for any *G, G*′ *∈* ℝ^*k×k*^,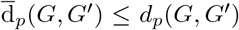. As *d*_*p*_ is a metric, for any *d*_*p*_(*G, G*) = 0, so 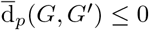. By definition, 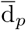 is non-negative, so 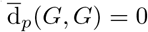.

Next, we check 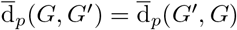. A simple check shows that for any *σ* ∈ `Perm` (*k*), we have that *d*_*p*_(*σ*(*G*), *G*′) = *d*_*p*_(*G, σ*^*−*1^(*G*′)). As *d*_*p*_ is a metric, we have that *d*_*p*_(*σ*(*G*′), *G*) = *d*_*p*_(*G, σ*(*G*′)). Putting the pieces together, for any bijection *σ*, we have that *d*_*p*_(*σ*(*G*), *G*′) = *d*_*p*_(*G, σ*^*−*1^(*G*′)) = *d*_*p*_(*σ*^*−*1^(*G*′), *G*). Hence, taking the infimum over all permutations,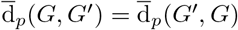.

Lastly, we check the triangle inequality. Let *G, G*′, *G*″ ∈ ℝ^*k×k*^. There are bijections *σ, σ*′∈ Perm(*k*) such that 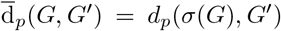 and 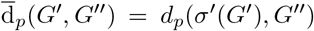. As in the symmetry case, *d*_*p*_(*σ*′(*G*′), *G*″) = *d*_*p*_(*G*′, *σ*^*′−*1^(*G*″)). By the triangle inequality for *d*_*p*_, we have that *d*_*p*_(*σ*(*G*), *G*″) + *d*_*p*_(*G*′, *σ*^*′−*1^(*G*″)) ≥ *d*_*p*_(*σ*(*G*), *σ*^*′−*1^(*G*″)). Again, as in the symmetry case, *d*_*p*_(*σ*(*G*), *σ*^*′−*1^(*G*″)) = *d*_*p*_(*σ*′(*σ*(*G*)), *G*″)), and finally, as *σ*′*σ* is a bijection, we have that 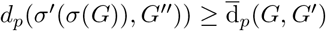. So, 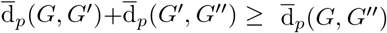, as required.

The case of 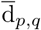 is a simple elaboration (in fact, the proof also works for 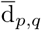, where the corresponding check that *d*_*p,q*_(*σ*(*G*), *G*′) = *d*_*p,q*_(*G, σ*^*−*1^(*G*′)) is more involved).

*Proof of Theorem 6.3*. The first part is as in the proof of Theorem 3.4, so we disprove the triangle inequality. We show that two sub-*L*-ecosystems of an *L*-ecosystem are distance 0 from the *L*-ecosystems, but can be distance *>* 0 from one another. Let *e ∈ ℰ*_*k*_ for some *k ∈* ℕ. Let *f* be a (strict) “sub-*L*-ecosystem” of *e*, i.e. *f ∈ ℰ*_*l*_ where *l < k* is downward-closed, and *G*^*f*^ is the restriction of *G*^*e*^ to *f*, so too for **x**^*f*^, *P* ^*f*^, and *c*^*f*^. Let *e −f* be the sub-*L*-ecosystem corresponding to “the rest” of *e*, i.e. *e−f ∈ ℰ*_|*k−l*|_, where 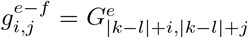, and so too for **x**^*e−f*^, *P* ^*e−f*^, and *c*^*e−f*^. Without loss of generality, we can choose *f* and *e−f* so that 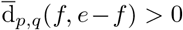. Then 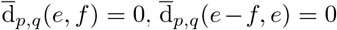, but 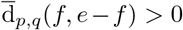.

For the last part, note that 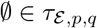 by definition, and that 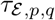 is trivially closed under unions.

*Proof of Theorem 6.5*. We consider only the case where *p, q ∈* ℝ, the *∞* cases are similar.

Suppose for a contradiction that there is a pair of disjoint sets, 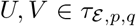, such that *U ∪ V* = *ℰ*. By the definition of 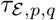, there are *e, r*_*e*_ and *f, r*_*f*_, with *r*_*e*_, *r*_*f*_ *>* 0 such that 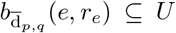 and 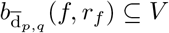.

We will construct a *g* ∈ *ℰ* such that 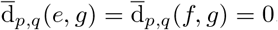. Without loss of generality, suppose *e* no more species than *f*. There are two cases.

Firstly, suppose *e* and *f* have some species in common, i.e. there is an *A* ⊆ |*e*| and a permutation *σ* : *A→* |*f* |such that the restriction, *R*, of the matrix *G*^*e*^ to the rows and columns *A*, is identical to restriction, *R*′, of the matrix *G*^*f*^ to the rows and columns of *f* (*A*) when those rows and columns are permuted according to *σ*. Let *g* be the induced sub-ecosystem of *e* including only the set of species *A*. Clearly, as ∈ *eℰ*, we have that ∈ *gℰ*. Then note that 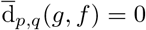 and 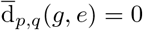. Hence *g ∈ U* ∩ *V*, contradicting that *U* and *V* are disjoint. So there is no such *U, V*.

Otherwise, suppose *e* and *f* have no species in common. Let *G*^*g*^ be the ‘gluing’ of *G*^*e*^ to *G*^*f*^, i.e. let *G*^*g*^ consist of the block *G*^*e*^, the block *G*^*f*^, and the remaining entries are taken to be 0 (cf. (Gluing), Section C). Adjoin the other data of *e* and *f* in the natural way to form a candidate model for |*e*| + |*f*|, which we call *g* (the matrix *P* ^*g*^ again is constructed as the block *P* ^*e*^, the block *P* ^*f*^, and the rest 0s; and *c*_*i,j*_ = 0 if *i* is ‘from’ *e* and *j* ‘from’ *f* and vice versa). It is a simple check that *g* satisfies each of the axioms A. to I. (inheriting them from *e* and *f*). Hence *g ∈ ℰ*. Note that, again, 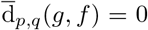 and 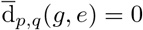. The proof continues as in the first case.

The last part follows from Proposition 3.1 and the fact that the only connected subsets of ℝ (with the standard topology) are the intervals.

## Footnotes

1 The production term is really the resultant term of production and (non-predation) death.

2 (Permuting) and (Shrinking) are *dependent* axioms in the sense that *G* is generated from (Basic Ecosystems) by (Gluing) alone.

## Notes

### Competing Interest Statement

The authors have declared no competing interest.

### Summary of Updates

Update some technical details in sections 3.2 and 5.2 for the case when different ecosystems have different numbers of species

